# Evolutionary genomics predicts probiotic persistence in corals

**DOI:** 10.1101/2025.06.16.659728

**Authors:** Mei Xie, Nan Xiang, Tianhua Liao, Chun Ting Cheung, Zhihao Xian, Pan Li, Chi Hin Lee, Wai Yin Tse, Kwong Kit Tsang, Congjuan Xu, Kaitlyn E. Ho, Qian He, Melanie Dörr, Hannah Manns, Xiaojun Wang, Danli Luo, Róisín Hayden, Emily Chei, Zhongyue Wan, Philip Thompson, Joseph Brennan, Raquel S. Peixoto, Guoxin Cui, Shelby E. McIlroy, Apple Pui Yi Chui, Christian R. Voolstra, Haiwei Luo

**Author notes:** These authors contributed equally to this work (Mei Xie, Nan Xiang, Tianhua Liao).

## Abstract

A universal bottleneck limiting probiotic efficacy in medicine, aquaculture, agriculture, and wildlife conservation is uncertain long-term colonization, necessitating repeated administration. We present an evolution-guided framework for probiotic identification based on genomic hallmarks of emerging host dependency, including widespread pseudogenization and insertion sequence proliferation driving genomic restructuring. Applied to coral reefs, we screened over 1,200 coral-associated bacterial isolates and identified *Ruegeria* MC10 as exhibiting these signatures. Its presence was associated with increased thermal tolerance of a model cnidarian. Following nursery application, MC10 persisted in reef corals for an 8-month monitoring period through a natural bleaching event, improving color retention and retaining algal photosynthesis performance. This work establishes a predictive, scalable pipeline for selecting persistent probiotics, directly addressing a central constraint on microbiome-based interventions across host systems.

## Main Text

Probiotics, defined as live microorganisms administered to confer health benefits, hold transformative potential across diverse fields from human medicine to wildlife conservation (*1–4*). A pervasive challenge limits their long-term efficacy and large-scale application: most administered microbes fail to establish stable, enduring colonization within host-associated microbiomes (*1, 2, 5–8*). This bottleneck necessitates repeated, costly, and often impractical reapplication, undermining the promise of probiotics as sustainable interventions.

Evolutionary genomics can provide a predictive solution to this colonization challenge. When bacteria transition from a free-living to a host-dependent lifestyle, their genomes undergo characteristic restructuring. Key hallmarks include the proliferation of insertion sequences (IS), mobile genetic elements that act as engines of genomic plasticity, coupled with the pseudogenization of core metabolic pathways (*9, 10*). These signatures reflect an evolutionary trajectory toward “host dependency”, where metabolic reliance can reinforce a persistent symbiotic association. We hypothesize that probiotic candidates selected *a priori* for these genomic features possess an intrinsic predisposition for long-term host residency.

Coral reefs present a critically threatened and functionally relevant system in which to test this predictive framework. These ecosystems sustain immense biodiversity and human livelihoods, yet they are collapsing under climate change (*11–13*). Since the 1950s, an estimated half of global coral cover has been lost (*14*). Even under the 1.5 °C Paris Agreement target, 70-90% of coral reefs are projected to be lost by 2050 (*15*). The breakdown of the coral-algal symbiosis during thermal stress triggers coral bleaching and subsequent mortality (*16*). The accelerating frequency of mass bleaching events, including the global events of 2014-2017 (*17*) and 2023-2025 (*18*), has left reefs with insufficient recovery intervals, necessitating active interventions to enhance resilience.

Here, we apply an evolutionary selection strategy to *Ruegeria*, a common bacterial genus member of coral holobionts (*19, 20*). From a collection of over 1,200 coral-associated bacterial isolates, we identified a *Ruegeria* population, MC10, exhibiting genomic signatures of early-stage host dependency. Using standardized acute heat-stress assays in the model cnidarian *Exaiptasia diaphana* H2, a companion study confirmed that exposure to MC10 was associated with increased thermal tolerance in a model cnidarian (*21*). This benefit was reduced or absent with sympatric *Ruegeria* strains isolated from the same coral colony and mucus compartment, which lack the genomic hallmarks of emerging host dependency that characterize MC10 (*21*). Following 4-week nursery inoculation with no subsequent reapplication in the field, we deployed MC10-and placebo-treated corals in an eight-month field trial encompassing a natural bleaching event happening in Hong Kong in the year 2024. We demonstrate that this candidate achieved sustained colonization *in hospite* that increased host thermal resilience. This work establishes a generalizable, predictive framework for selecting enduring probiotics, offering a solution to a universal probiotic limitation and providing a timely, scalable tool for ecosystem resilience.

## Results

### Ruegeria population MC10 as a putative next-generation coral probiotic

To identify candidate probiotics exhibiting genomic hallmarks of host dependency, we cultured 419 *Ruegeria* isolates from five coral species (*Acropora* cf. *samoensis*, *Acropora solitaryensis*, *Favites pentagona*, *Platygyra acuta*, and *Oulastrea crispata*) collected across nine sites spanning environmental gradients in Hong Kong (**Fig. S1**). Bacterial isolates were primarily obtained from coral mucus and skeleton (**Fig. 1A**), sampling from potentially transient surface associates to more persistent tissue-and skeleton-associated taxa.

**Fig. 1.**
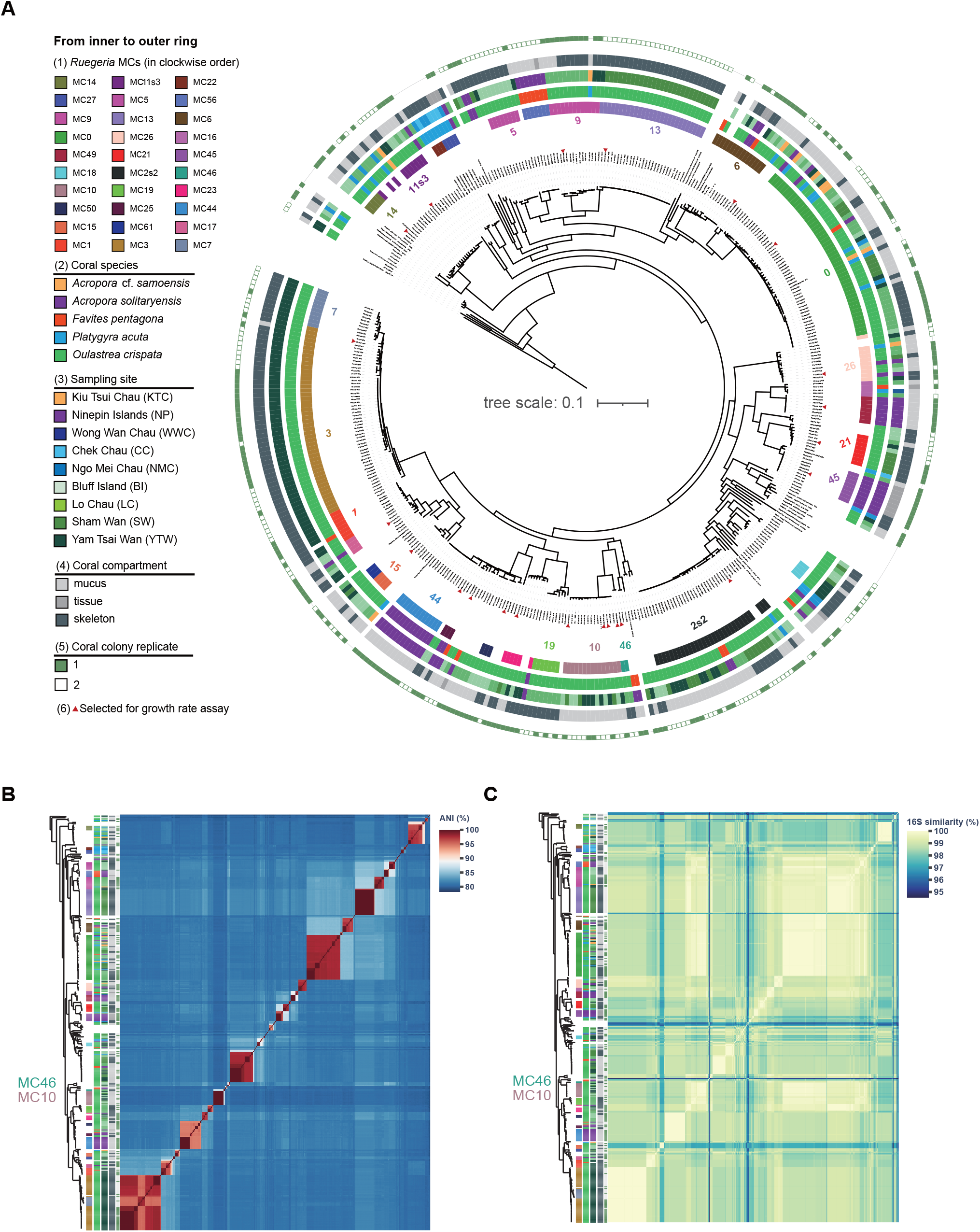
***Ruegeria* MC10’s phylogenetic position. (A)** Rooted maximum-likelihood phylogenomic tree comprising 419 *Ruegeria* isolates associated with five coral species sampled from nine Hong Kong reef sites, along with 26 *Ruegeria* genomes retrieved from public databases. This tree was constructed with IQ-TREE based on concatenated single-copy orthologous genes at the amino acid level. PopCOGenT was used to delineate the *Ruegeria* populations (i.e., main clusters or MCs). The 19 MCs that contain representative members additionally sequenced by Nanopore were labelled with numbers in the tree. From the inner to outer rings: (1) denotes 29 *Ruegeria* populations which consists of at least three isolates as defined by PopCOGenT, along with MC46, the sister group of MC10; (2) - (5) denote the source of *Ruegeria* isolates with each corresponding to coral species, sampling site, coral compartment, and coral colony replicate, respectively. The heatmaps of pairwise (**B**) whole-genome average nucleotide identity (ANI) and (**C**) 16S rRNA gene identity among 419 *Ruegeria* isolates. From left to right columns: (1) denotes the 30 *Ruegeria* populations with the same marker colors and names as in Fig. 1A; (2) - (5) denote the source information for *Ruegeria* isolates, including coral species, sampling locations, coral compartments, and coral colony replicates.

Using PopCOGenT, a method that delineates populations (main clusters, MCs) based on recombination barriers without phylogenetic tree input (*22*), we resolved 116 genetically cohesive *Ruegeria* populations, 29 of which contained at least 3 isolates (*21, 23*). Phylogenomic analysis confirmed monophyly for most populations, validating PopCOGenT’s accuracy for tracking probiotic diversity at an ecologically meaningful unit (**Fig. 1A**). Within a population, average nucleotide identity (ANI) typically exceeded 95%, whereas between populations, ANI fell below 90% **(Fig. 1B**). By contrast, 16S rRNA gene identity did not reliably differentiate population-level differences, as even distantly related populations can share >99% sequence similarity (**Fig. 1C**).

To identify genomic signatures of host dependency, we additionally sequenced and assembled 34 closed genomes spanning the *Ruegeria* phylogenetic diversity (**Fig. 2 & Fig. S2**). This analysis revealed that population MC10 exhibits the genomic hallmarks of evolutionary recent endosymbionts, specifically insertion sequence (IS) expansion (**Fig. 2A**) and pseudogene proliferation (**Fig. 2B & Fig. S3**). These signatures, which reflect an ongoing transition toward host dependency, were absent or minimal in other populations, making MC10 a prime candidate for probiotic evaluation. Within MC10, frameshift and nonsense mutations disrupted genes across multiple essential metabolic pathways. Disruptions in the tricarboxylic acid (TCA) cycle were prominent, affecting genes including aconitate hydratase (*ACO*), isocitrate dehydrogenase (*IDH1*), succinyl-CoA synthetase (*sucC*), malate dehydrogenase (*mdh*), and succinate dehydrogenase (*sdhA*) (**Fig. 3A & 3B**). The location of these mutations (**Fig. 3C**) suggests a gradient of functional impact: 5’-terminal disruptions are expected to severely impair protein function, whereas mutations nearer the 3’ terminus may allow for partially functional gene products. This spectrum of mutational effects likely underpins the retained, albeit reduced, metabolic capacity enabling MC10’s culturability. Pseudogenization was not restricted to the TCA cycle but was widespread across glycolysis, the pentose phosphate pathway, gluconeogenesis, amino acid biosynthesis, and vitamin and cofactor synthesis pathways (**Fig. 3D**). These pseudogenized loci typically lacked functional paralogs, suggesting a heightened metabolic reliance on the coral host.

**Fig. 2.**
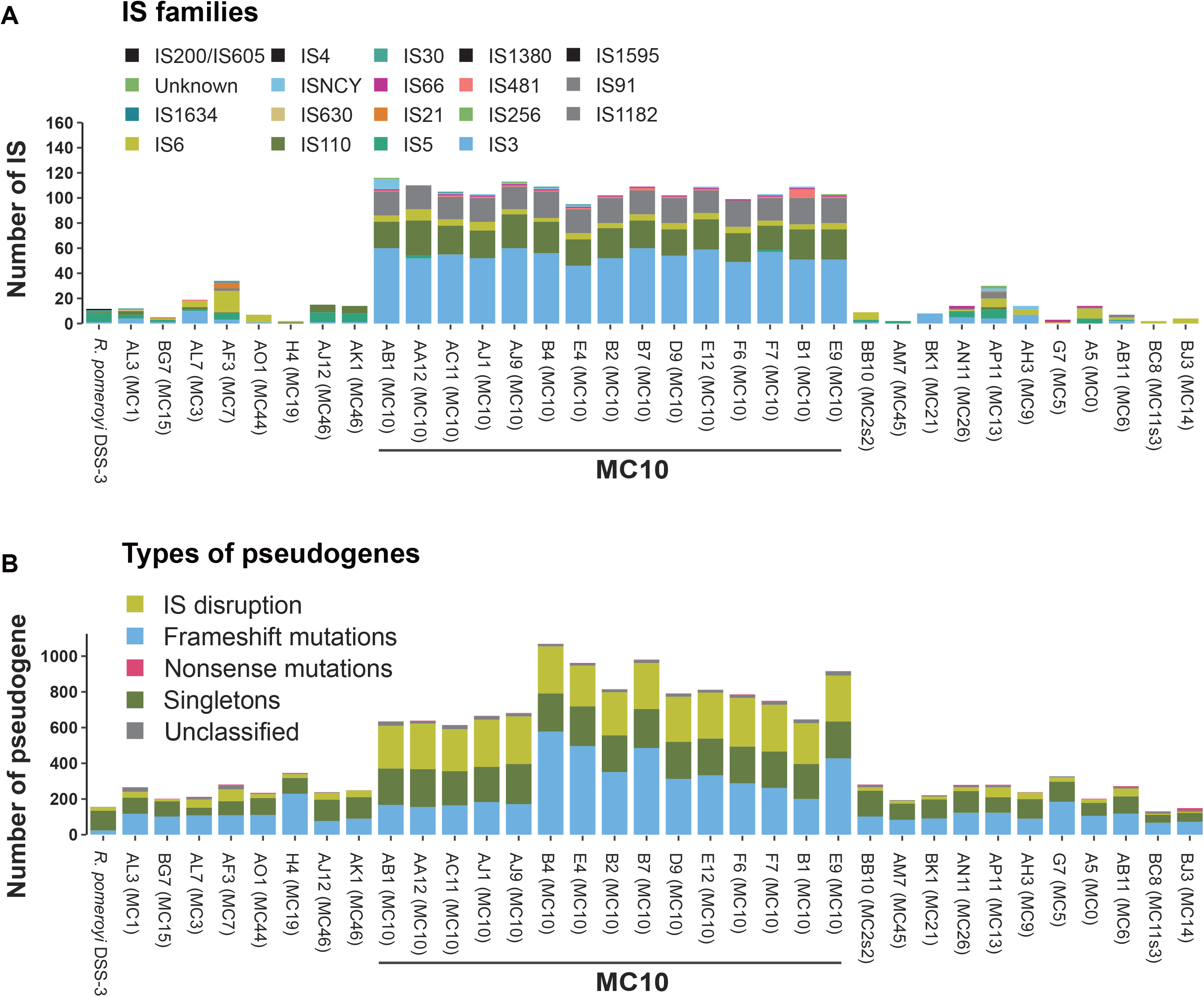
***Ruegeria* MC10’s genomic signatures showing recent host dependency. (A)** Stacked bars showing the number of IS and IS family composition in 34 newly closed *Ruegeria* genomes and the reference genome *Ruegeria pomeroyi* DSS-3. **(B)** Number of pseudogenes in 34 newly closed *Ruegeria* genomes and the reference genome *R. pomeroyi* DSS-3. Pseudogenes were grouped by the mechanism of formation, namely “IS disruption”, “Frameshift mutations”, “Nonsense mutations”, and “Singletons”. “Unclassified” pseudogenes are those that cannot be assigned to any of the three defined mechanisms.

**Fig. 3.**
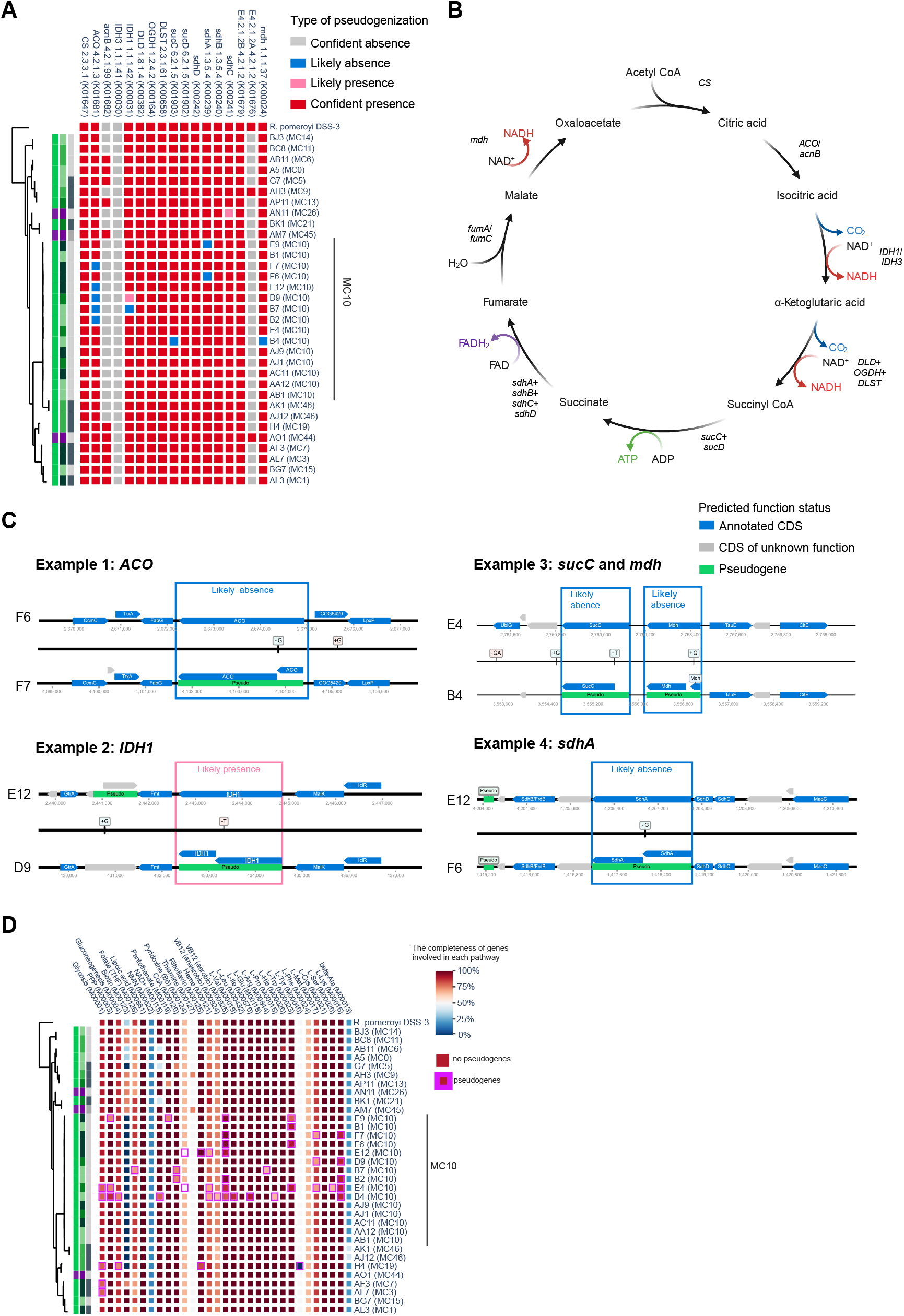
Pseudogenes of MC10 were identified in genes involved in tricarboxylic acid (TCA) cycle and other essential metabolic pathways. **(A)** TCA cycle-associated pseudogenes were classified into four categories, including “Confident presence”, “Confident absence”, “Likely presence”, and “Likely absence”. **(B)** The schematic representation and corresponding gene names responsible for the TCA cycle are derived from the KEGG database. Alternative genes are denoted with a slash (e.g., “IDH1/IDH3”), while component genes (subunits of a complex) are indicated by a plus sign (e.g., “*sucC* + *sucD*”). **(C)** TCA cycle-associated pseudogenization in MC10 due to frameshift mutations was exemplified by five genes: aconitate hydratase (ACO), isocitrate dehydrogenase (IDH1), succinyl-CoA synthetase (*sucC*), malate dehydrogenase (*mdh*), and succinate dehydrogenase (*sdhA*). These examples show the comparison of intact genes (top) versus pseudogenes (bottom). Annotated coding sequences (CDS) were shown in blue, with CDS of unknown function shown in grey, pseudogenes shown in green, and frameshift-inducing indels indicated (e.g., “+G” means guanine insertion). **(D)** Pseudogenes of MC10 identified some genes responsible for essential metabolic pathways, including glycolysis, pentose phosphate pathway, gluconeogenesis, amino acid synthesis, vitamin and cofactor synthesis. The color of blocks represents the completeness of genes involved in different pathways for each *Ruegeria* isolate, with a purple frame indicating the presence of pseudogenes.

Within MC10, we observed functional heterogeneity in these hallmark signatures. Some strains (e.g., MC10-B4) exhibited extensive pseudogenization across the pathways described, whereas others (e.g., MC10-AJ1) retained largely complete pathways (**Fig. 3A & 3D**). This within-population variation presents a potential source of difference in strain-level efficacy. MC10 collectively exemplifies the defining genomic features of a next-generation probiotic candidate: pronounced IS-driven genomic restructuring coupled with expanded pseudogenization (**Fig. 2A and 2B**), hallmarks of a lineage transitioning toward host dependency.

Phylogenetically, MC10’s closest relative is MC46 (**Fig. 1**). Although the two populations shares 99.45% 16S rRNA gene identity, they diverged genomically, with an ANI of 86.88 ± 0.04% (**Fig. 1B**). This ANI falls well below the 95% within-population threshold and is consistent with the between-population expectation, confirming that MC10 and MC46 are distinct populations despite their high 16S similarity (**Fig. 1C**). In contrast to MC10, MC46 lacks IS expansion and widespread pseudogenization (**Fig. 2**). This contrast underscores MC10’s distinct genomic trajectory toward host association.

### Reduced free-living growth distinguishes Ruegeria MC10 from related populations

The genomic hallmarks of early-stage host dependency prompted us to investigate whether this evolutionary trajectory is associated with a physiological cost to free-living growth. We performed a phylogenetically-informed comparison of maximum growth rates (μ_max_) across a diverse panel of 24 *Ruegeria* isolates (**Fig. 1A**). This panel included three strains from population MC10, representing its spectrum of pseudogenization from the extensively pseudogenized MC10-B4 to isolates (e.g., MC10-AJ1) with minimal pseudogenization in essential pathways, all sampled from distinct reef sites; two strains from its sister population MC46; and 18 phylogenetically diverse strains from 18 other *Ruegeria* MCs spanning various hosts, compartments, and sites, plus the free-living model *R. pomeroyi* DSS-3 (*24*).

Consistent with predicted metabolic constraints arising from pseudogenization, members of population MC10 exhibited significantly lower maximum growth rates compared to all other *Ruegeria* populations under standardized laboratory conditions (Phylogenetic ANOVA, Brownian motion model, *P* = 0.009; **Fig. 4A & 4B**; **Table S1**). This reduction was particularly evident when comparing MC10 to its closest relative, MC46, which lacks IS expansion and pseudogene proliferation (**Fig. 2**). MC46 strains displayed substantially higher growth rates, underscoring that the attenuated growth is a specific phenotypic correlate of MC10’s unique genomic trajectory toward host association, rather than a shared trait of closely related lineages.

**Fig. 4.**
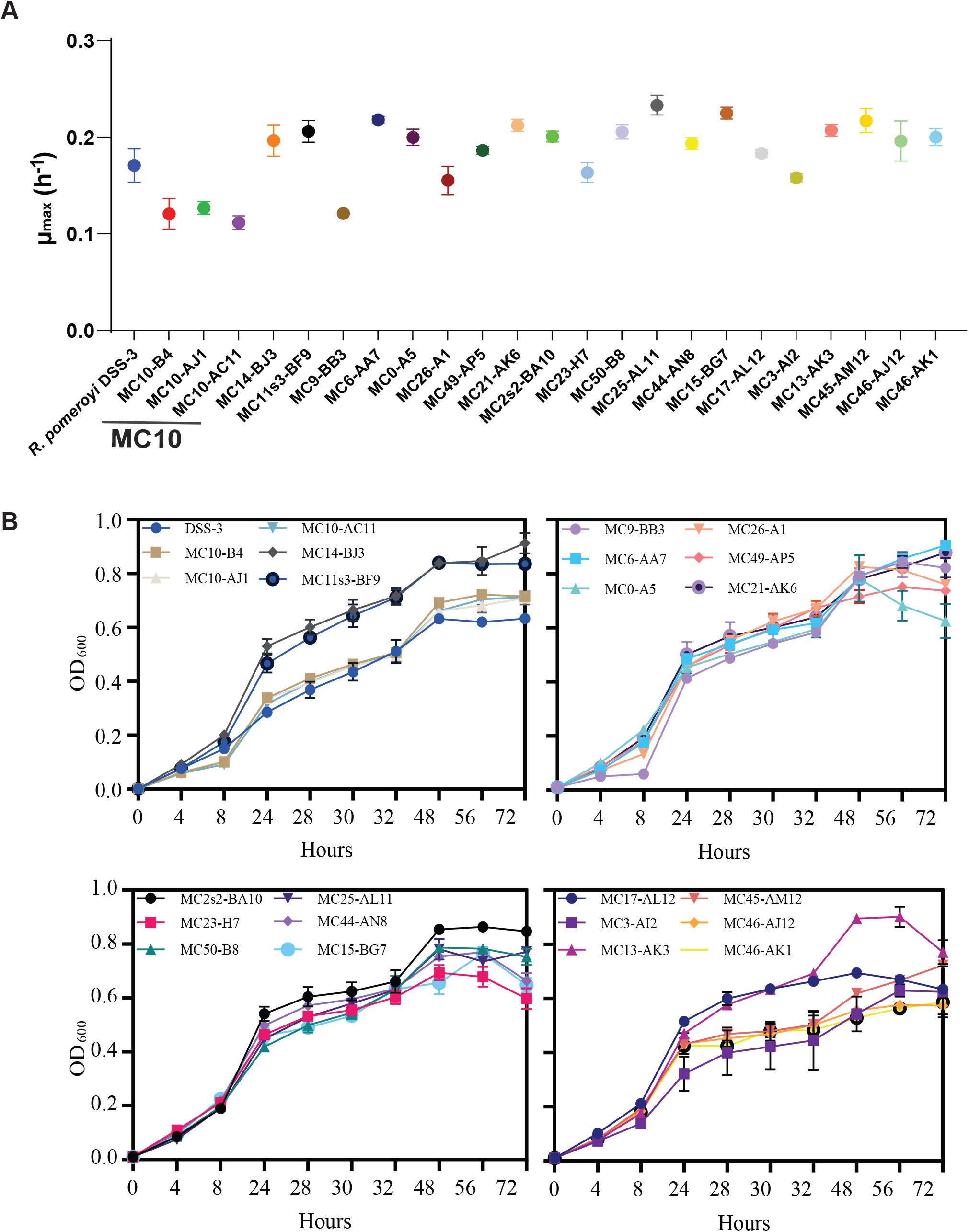
***Ruegeria* MC10’s free-living physiological cost. (A)** Maximum growth rate (μ_max_, h^-1^) comparison. Bars represent the mean μ_max_ ± standard deviation of the MC10 population, the non-MC10 *Ruegeria* population and a free-living model *R. pomeroyi* DSS-3. Growth rates were calculated from the exponential phase of growth curves. Significant difference (PhylANOVA with Brownian motion evolution model, *P* = 0.009) was observed in growth rates between MC10 versus non-MC10 counterparts. **(B)** Growth curves of the 24 *Ruegeria* isolates, including 3 strains from the MC10 population, 20 strains from the non-MC10 population and the model strain *Ruegeria pomeroyi* DSS-3, cultured in Marine Broth 2216 with shaking at 200 rpm and 28°C. Data were shown as mean ± standard deviation (SD) (n = 3 replicates each).

Despite this constraint on growth rate, MC10 remains fully culturable and achieves a robust cell yield. The representative strain MC10-B4 (μ_max_ = 0.12 h^-1^) reached a stationary-phase density of 9.92 ± 0.14 Log_10_ CFU/mL, which was significantly higher than the yields of the sympatric control strain MC15-BG7 (9.44 ± 0.03 Log_10_ CFU/mL) and the model *R. pomeroyi* DSS-3 (9.25 ± 0.10 Log_10_ CFU/mL).

### Ruegeria MC10 is present across diverse coral species

To assess the ecological distribution and host associations of *Ruegeria* MC10, we performed targeted amplicon sequencing of two population-resolving marker genes, *parC* (encoding a topoisomerase IV subunit) and *ATP5B* (encoding an ATP synthase subunit). These markers were selected from over 700 conserved single-copy *Ruegeria* gene families. Their ability to resolve PopCOGenT-defined populations into monophyletic groups was validated in a previous study (*23*). Custom primers achieved high recovery rates (64.8% for *parC*; 74.3% for *ATP5B***; Table S2**), enabling sensitive detection across samples.

Although pure cultures of MC10 were originally sourced from *Oulastrea crispata* (**Fig. 1A**), culture-independent amplicon data revealed its presence in all five taxonomically distinct coral species (*Oulastrea crispata*, *Acropora solitaryensis*, *Plesiastrea versipora*, *Porites* sp., and *Platygyra acuta*) sampled from diverse Hong Kong reef sites, as well as in seawater and sediment samples (**Fig. 5**). MC10 detection varied among colonies of the same species. Where present, MC10 typically constituted <2% of local coral-associated total *Ruegeria* communities. Estimates of its relative abundance based on *parC* and *ATP5B* amplicons showed no linear correlation at low proportions (<0.5% of total *Ruegeria*; **Fig. S4**), reflecting methodological challenges in quantifying low-abundance populations. We also attempted to quantify the absolute MC10 abundance via qPCR but did not obtain reliable signals (**Fig. S5**).

**Fig. 5.**
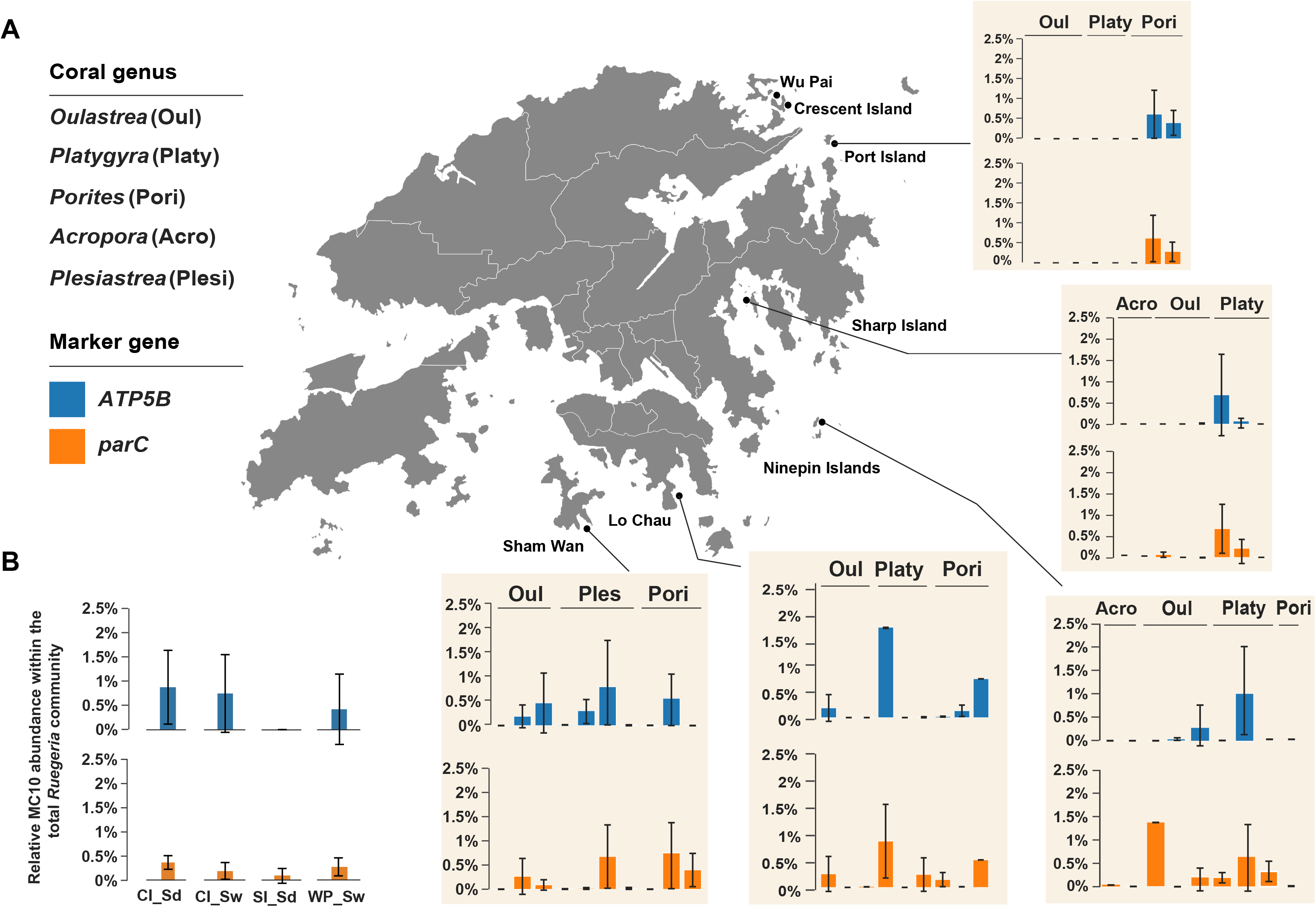
Relative abundance of *Ruegeria* MC10 in coral holobionts and environmental samples from Hong Kong reefs. **(A)** The relative abundance of MC10 in *Ruegeria* communities was quantified based on population-resolving *ATP5B* and *parC* amplicon sequencing across natural reefs in Hong Kong. Different bar plots under the same coral genus represented different coral colonies (i.e., 15 colonies for *Oulastrea* sp., 11 for *Platygyra* sp., 10 colonies for *Porites* sp., 4 colonies for *Acropora* sp., and 4 colonies for *Plesiastrea* sp. across five sites). Each bar plot was shown as mean ± standard deviation (SD) calculated from three compartments (i.e., mucus, tissue, and skeleton) for the same coral colony. **(B)** The relative abundance of MC10 in seawater and sediment samples across different locations. The error bar denotes standard deviation among three replicates. CI, Crescent Island; SI, Sharp Island; WP, Wu Pai. Sd, sediment; Sw, seawater.

### Sustained association and thermal tolerance in reef corals following Ruegeria MC10 application

Prior to field deployment, we assessed the colonization capacity predicted by MC10’s evolutionary genomics profile. In a controlled *ex situ* assay, nubbins derived from multiple *Acropora pruinosa* colonies (with fragments from each colony distributed across treatments) received twice-weekly inoculations over four weeks with strain MC10-B4, the sympatric control strain MC15-BG7 (isolated from the same coral colony and compartment), or an autoclaved filtered seawater (AFSW) placebo. Amplicon sequencing revealed that MC10-B4 achieved a median relative abundance of 18.331% within the coral *Ruegeria* community (0.003% of total bacteria), significantly higher than the placebo (2.410% of *Ruegeria*, <0.001% total bacteria; *P* = 0.002 and *P* = 0.007, respectively; one-way ANOVA; **Fig. 6A**, **Table S3**). In contrast, corals inoculated with MC15-BG7 showed no significant increase in MC15 abundance (median 3.242% of *Ruegeria*, <0.001% total bacteria) over the placebo control (0.251% of *Ruegeria*, <0.001% total bacteria; *P* = 0.136 and 0.281; **Fig. 6B**, **Table S3**).

**Fig. 6.**
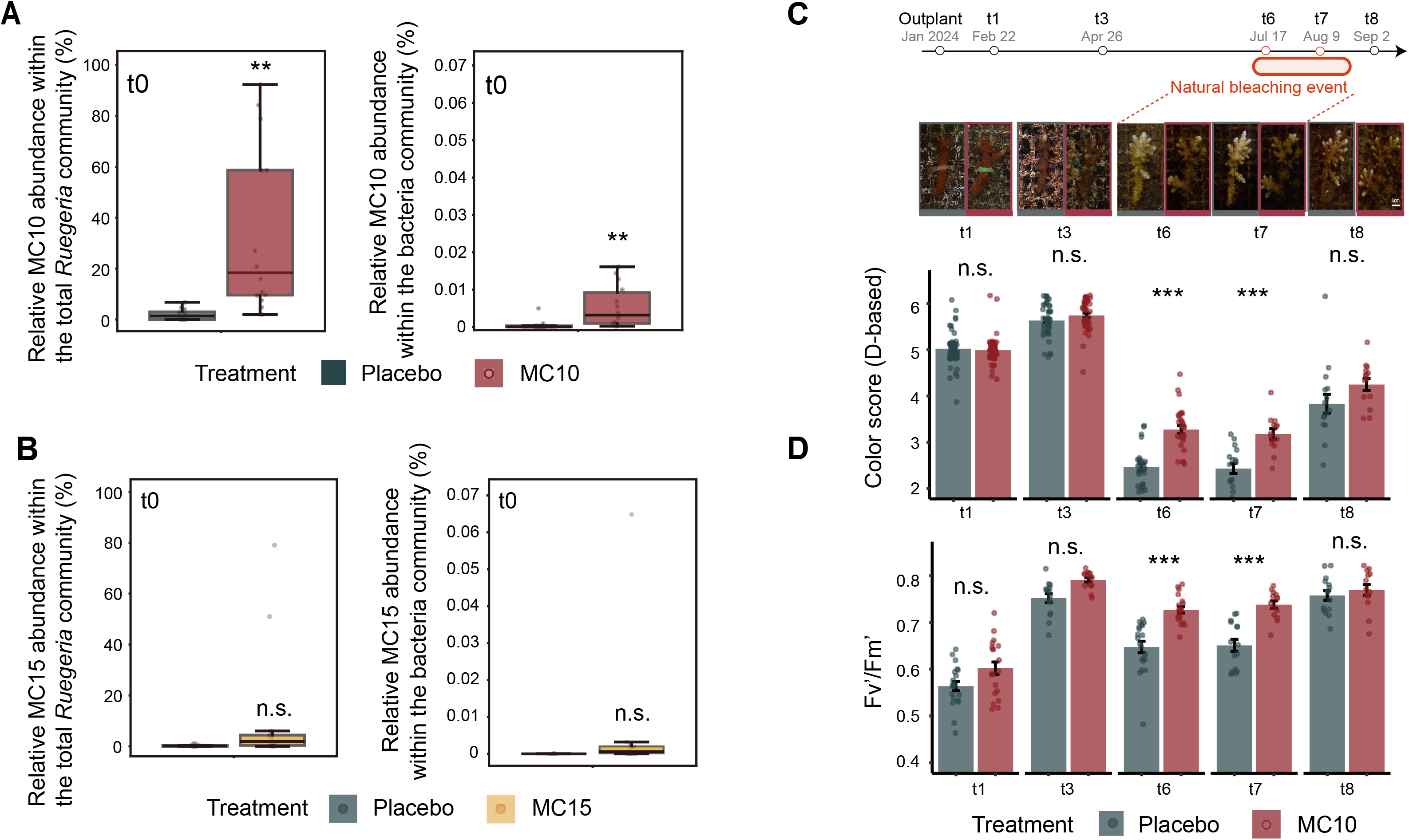
Application of *Ruegeria* MC10-B4 through field trials on coral *Acropora pruinosa*. The relative abundance of **(A)** MC10 and **(B)** MC15 in the *Ruegeria* (left axis; based on *parC* amplicon) and total bacteria community (right axis; combined 16S rRNA and *parC* gene amplicon) following twice-weekly inoculations over a month. The central line of each box indicates the median value, with the upper and lower error bars showing maximum and minimum potential variabilities, respectively. **(C)** Coral phenotype and coloration, and **(D)** effective quantum yield (F_v_′/F_m_′) were assessed on MC10-B4-inoculated and placebo-treated corals across five timepoints: 1, 3, 6, 7 and 8 months post-outplanting (t1, t3, t6, t7, t8, respectively). Data were presented with the mean values and standard errors.

Following this *ex situ* validation, MC10-B4-inoculated and placebo-treated *A. pruinosa* fragments were outplanted to adjacent reef sites (40 m apart) in January 2024 (**Fig. S6**). Corals were monitored over eight months at defined timepoints: after nursery inoculation but before outplanting (**t0**), and at 1, 3, 6, 7, and 8 months post-outplanting (**t1**, **t3**, **t6**, **t7**, **t8**). A natural, region-wide bleaching event occurred in Hong Kong between July and September 2024, during which degree heating weeks (DHW) reached 8.8-18 °C-weeks (**Fig. S7**), exceeding established NOAA bleaching thresholds.

MC10-B4-inoculated corals exhibited enhanced thermal tolerance specifically during the natural bleaching event. At the peak of thermal stress (**t6** and **t7**), probiotic-treated corals showed significantly reduced bleaching severity compared to placebo controls (CoralWatch Coral Health Chart coloration scores at **t6**: 3.3 ± 0.5 vs. 2.5 ± 0.4, *P* < 0.001; **t7**: 3.2 ± 0.4 vs. 2.4 ± 0.4, *P* < 0.001; two-way ANOVA, Tukey’s HSD; **Fig. 6C, Table S4**). The operational efficiency of PSII under ambient light conditions (F_v_′/F_m_′) was also significantly higher in the probiotic group during this period (**t6**: *P* < 0.001; **t7**: *P* < 0.001; two-way ANOVA, Tukey’s HSD; **Fig. 6D, Table S5**). A significant treatment-by-time interaction (*P* < 0.001, two-way ANOVA, interaction effect) indicated that the probiotic’s effect was contingent on the presence of thermal stress. Following the thermal stress peak but with DHW levels still elevated above stress-relief thresholds at **t8**, both groups exhibited partial recoloration.

To investigate the mechanism underlying the observed thermal resilience, we conducted a control experiment to exclude potential effects from nutritional supplementation, transient priming by live cells, or postbiotic contributions from bacterial components and metabolites. *Acropora pruinosa* nubbins were exposed for 28 days to live, autoclaved, or pasteurized cells of MC10-B4 and MC15-BG7, using the same twice-weekly inoculation scheme applied in the nursery. An AFSW placebo control was included, yielding seven experimental groups in total. A holistic physiological assessment showed that live inoculation with either bacterial strain did not alter the coral’s baseline state across key metrics, including dark-adapted maximum photosynthetic efficiency (F_v_/F_m_) (**Fig. 7A**), coloration (**Fig. 7B**), host protein content (per surface area) (**Fig. 7C**), total lipid energy content (per dry weight) (**Fig. 7D**), chlorophyll a concentration (per surface area) (**Fig. 7E**), symbiont density (per surface area) (**Fig. 7F**), and host fluorescent protein content (Relative Fluorescence Units per host protein weight) (**Fig. 7G**), compared to the placebo control (two-way ANOVA, Tukey’s HSD, *P* > 0.05; **Table S6**). Isolated changes were observed in fragments exposed to pasteurized MC10-B4 cells, which exhibited a reduction in F_v_/F_m_ (**Fig. 7A**) and an increase in host fluorescent protein content (**Fig. 7G**) compared to placebo controls (*P* = 0.004 and 0.001, respectively). In contrast, fragments exposed to pasteurized MC15-BG7 showed a significant increase in total lipid energy content (*P* = 0.003) (**Fig. 7D**). These effects were not observed with pasteurized counterparts of the other strain or with autoclave-killed cells of either strain.

**Fig. 7.**
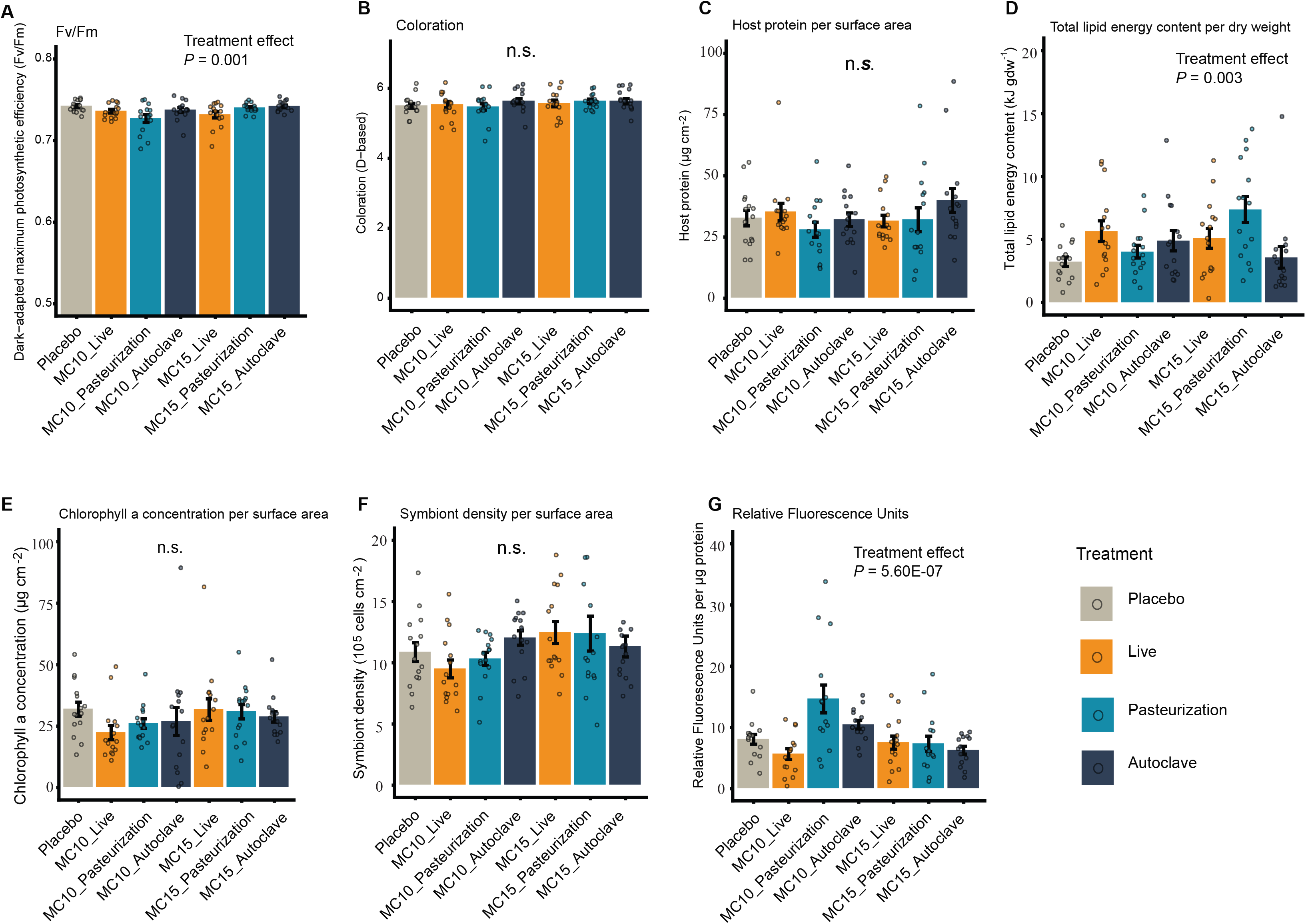
Physiological metrics of coral holobionts after one-month inoculation confirming equal healthy coral physiological status without heterotrophic feeding benefit before the outplanting. **(A)** F_v_/F_m_. **(B)** Coloration. **(C)** Host protein per surface area. **(D)** Total lipid energy content. **(E)** Chlorophyll a concentration per surface area. **(F)** Symbiont density per surface area. **(G)** Relative fluorescent protein unit. All physiological data were presented in barplots with mean and standard errors (n = 15). Scatter points show raw data. Statistical analysis was done across treatments (one-way ANOVA, Tukey HSD, *P* < 0.05).

To visually confirm the persistent presence of MC10 within host tissues, we used fluorescence *in situ* hybridization (FISH) with a population-specific probe. Because conventional 16S rRNA probes lack the resolution to distinguish MC10 from other *Ruegeria* populations, we developed a 215 bp probe targeting the single-copy *parC* gene. Probe specificity was validated using pure cultures of MC10-B4 (**Fig. 8A**). In coral samples collected immediately after nursery inoculation (**t0**) and at six (**t6**) and eight (**t8**) months post-outplanting, the probe revealed MC10 cells localized within the coral gastrodermis (**Fig. 8B**). Specificity in holobiont samples was confirmed using a nonsense probe control (**Fig. 8B**). MC10 was not necessarily co-localized with Symbiodiniaceae. The low natural abundance of MC10 within *Ruegeria* communities (median 0.23%) and of *Ruegeria* within total coral microbiomes (median 0.02%) precluded precise quantification via amplicon sequencing, falling below typical detection threshold (0.5%) for reliable amplicon sequence variant (ASV) resolution (*25*).

**Fig. 8.**
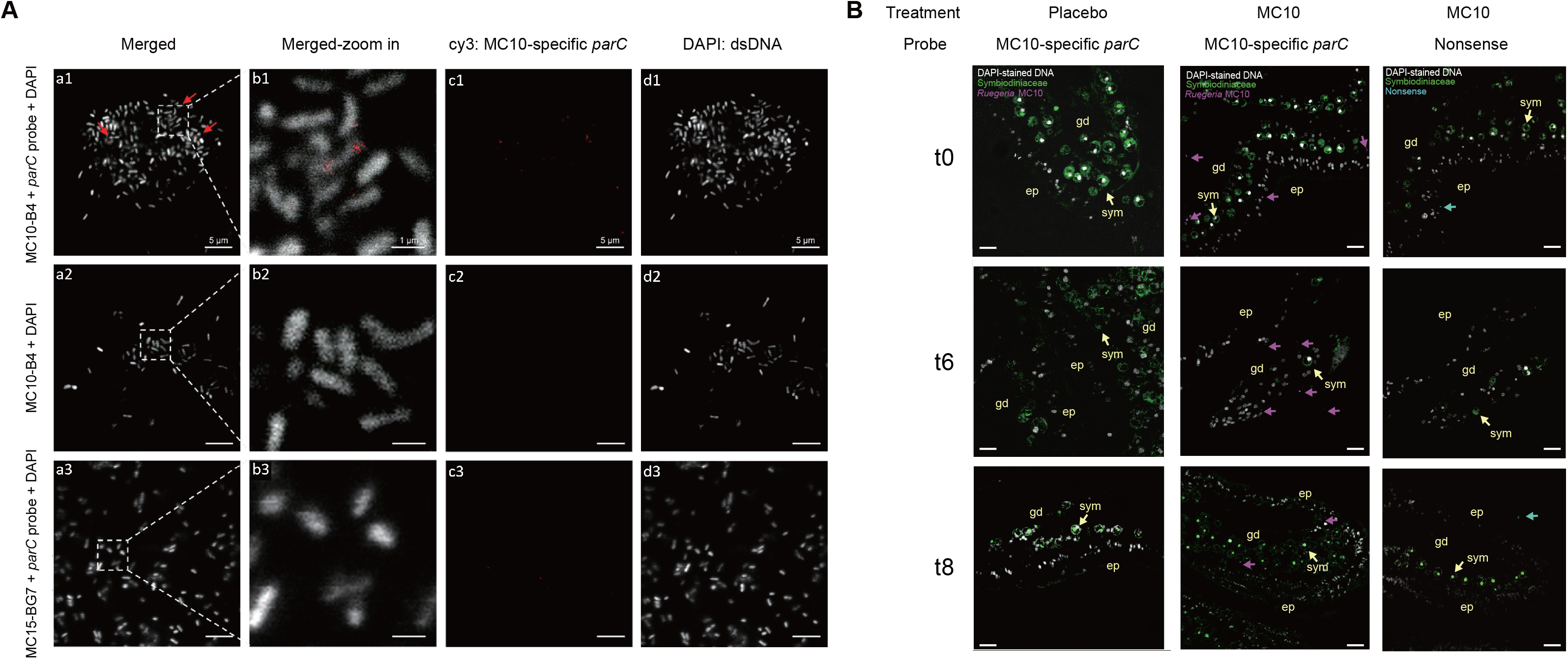
Validation of the MC10-specific *parC* probe and tracking of *Ruegeria* MC10 within the coral tissue using Fluorescence *in situ* hybridization (FISH). (A) The MC10-specific *parC* FISH probe selectively targets *Ruegeria* strain MC10-B4. Confocal microscopy images of (1) *Ruegeria* MC10-B4 hybridized with the Cy3-labeled *parC* probe, (2) MC10-B4 without probe (blank control), and (3) non-target strain MC15-BG7 hybridized with the same probe. For each row, panels show: (a) merged image, (b) zoom of the region indicated by the dashed boxes in (a), (c) Cy3 channel (probe signal), and (d) DAPI channel (dsDNA stain). Red arrows indicated the probe signals co-localized with DAPI-stained MC10-B4 cells. All images were acquired under identical settings (excitation wavelength, laser power, detector gain, and detection range). Scale bars: a, c, and d: 5 μm; b: 1 μm. **(B)** Localization of MC10 in coral *A. pruinosa* after one-month inoculations (t0, before outplanting), six months post-outplanting (t6, during bleaching), and eight months post-outplanting (t8, post-bleaching). Representative confocal images depicting the hybridization of *parC* probe specifically targeting MC10 labeled with Cy3 (purple). The purple arrows pointed at the specific locations of MC10 in the coral gastrodermis. The blue arrows pointed at the non-specific binding sites with an EUB nonsense probe. Grey, DAPI-stained dsDNA; Green, autofluorescence of Symbiodiniaceae; Scale bars, 10 μm.

Bacterial community dynamics in outplanted corals were assessed via *parC* and 16S rRNA gene amplicon sequencing at multiple timepoints (**t0, t1, t3, t6**, and **t8**; **t7** samples unavailable). Analysis of β-diversity revealed that bacterial communities in MC10-B4-inoculated corals became progressively more similar to those in placebo corals over the monitoring period (**Fig. 9A**; ANOSIM). During the peak bleaching phase (**t6**), genus-level analysis identified a significant enrichment of several Bacteroidetes genera (*Cryomorpha*, *Spirosoma*, *Sunxiuqinia*, *Gelatiniphilus*, *Aureispira*, and *Flavobacterium*) and a depletion of multiple Roseobacteraceae genera (*Sulfitobacter*, *Roseovarius*, *Pseudoruegeria*, *Donghicola*, *Aliiroseovarius*, and *Phaeobacter*) in MC10-B4-inoculated corals compared to the placebo group (FDR-corrected *P* < 0.05; ANCOM-BC; **Fig. 9B**). Most of these taxa showed no differential abundance during pre-bleaching (**t3**) or post-bleaching (**t8**) phases (**Fig. S8**), indicating that MC10 inoculation was associated with a specific restructuring of the coral microbiome during thermal stress.

**Fig. 9.**
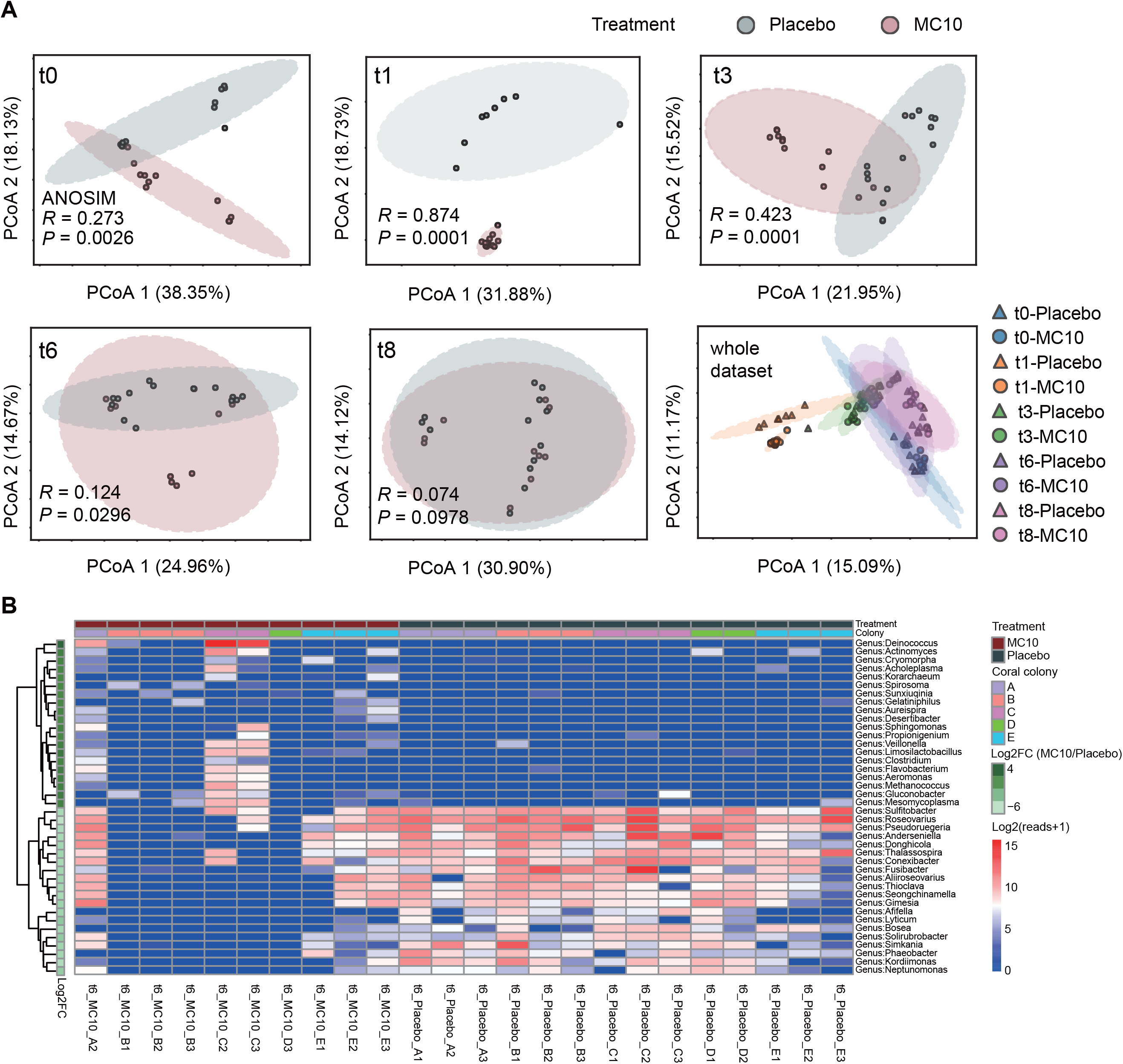
Coral microbiome dynamics during the 8-month field trial. **(A)** Principal Coordinates Analysis (PCoA) of coral holobiont-associated bacterial communities based on Bray-Curtis dissimilarity, comparing placebo and MC10-B4-treated groups across five timepoints: baseline (t0, pre-outplanting) and 1, 3, 6, and 8 months post-outplanting (t1, t3, t6, t8, respectively). Ellipses represent 95% confidence intervals for group centroids. ANOSIM was used to assess the separation of bacterial communities between placebo and MC10-B4-treated corals, with R values interpreted as: little to no (R < 0.25), weak (0.25 ≤ R < 0.5), moderate (0.5 ≤ R < 0.75), and strong (R ≥ 0.75). **(B)** Differentially abundant taxa between placebo and MC10-B4-treated corals at t6 (during bleaching) identified by ANCOM-BC analysis (*P* < 0.05, FDR corrected). A heatmap displaying top 20 significantly upregulated (positive log_2_FC) and downregulated (negative log_2_FC) genera, respectively. A log_2_-transformed read counts (log_2_[reads+1]) were shown to indicate the absolute fold-change magnitudes of taxa. Pseudocounts were added to handle the case of zeros in the calculation.

## Discussion

The accelerating loss of biodiversity under climate change demands innovative strategies to enhance ecosystem resilience. Probiotics have shown promise, yet their long-term efficacy is often limited by transient colonization, a fundamental limitation on scalable *in situ* application (*1, 2, 5–8*). Here, we propose and validate a framework to overcome this challenge: the selection of probiotic candidates based on evolutionary genomic signatures indicative of a transition towards host dependency, which predicts stable host association. We demonstrate this framework in the context of coral reefs. Focusing on the coral-associated bacterial genus *Ruegeria* (*19, 20*), we identified population MC10, which exhibits hallmarks of early-stage host dependency. A nursery inoculation enabled MC10 to be retained in coral tissues throughout an 8-month field trial, where its persistent colonization conferred increased host thermal resilience during a natural bleaching event. This work establishes a predictive probiotic selection pipeline that combines evolutionary trajectory with functional validation.

### Evolutionary genomic signatures as predictors of probiotic persistence

Common approaches to selecting marine probiotics have primarily relied on screening for functional traits such as pathogen inhibition, ROS degradation, or nutrient provisioning (*26, 27*), with the hope that selected strains will persist in the host. A key limitation for scalable application, however, is the uncertainty regarding their long-term residency within host tissues (*28*). Our framework inverts this process: we first select for evolutionary signatures predictive of stable colonization, and then empirically test whether the persistent strain provides a host benefit.

This strategy is grounded in the hypothesis that specific genomic signatures predict a bacterial lineage’s propensity for stable host association. In MC10, we observed hallmarks of genomic restructuring and decay, including IS expansion and pseudogene proliferation. IS proliferation serves as an engine of genomic plasticity, facilitating deletions and rearrangements that accelerate host adaptation (*29, 30*). This state enables the fixation of conditionally deleterious mutations, such as those disrupting core metabolism in MC10. Such genomic restructuring incurs a defined phenotypic cost: MC10 exhibited a significantly reduced maximum growth rate under free-living conditions compared to related *Ruegeria* populations. This attenuated independent proliferation capacity phenotypically validates the genomic trajectory toward host dependency. The resulting metabolic reliance reinforces symbiosis, transforming a physiological cost in one environment into a stabilizing mechanism in another.

The genomic signatures observed in *Ruegeria* MC10 mirror those in terrestrial model symbionts such as *Sodalis glossinidius*, where they signify a transitional state toward host dependency (*9, 10*). Proactively applying these evolutionary principles to select probiotics represents a distinct strategy for conservation. Our study tests the key functional implication: that this genomic trajectory facilitates stable, long-term association. Supporting this, a companion trait study (*21*) reveals that MC10’s genomic architecture is coupled with a functional profile conducive to host residency, including robust biofilm formation and host-induced “motility-to-sessility” proteomic reprogramming, responses not observed in sympatric *Ruegeria* isolates like MC15-BG7. The sustained colonization and enhanced thermal resilience demonstrated by *Ruegeria* MC10 confirm the feasibility of this evolution-guided approach.

### Validating the framework: From laboratory predictions to field trial design

A primary objective of this work was to introduce and provide a field validation for a two-part selection framework. This framework first employs evolutionary genomics to identify probiotic candidates with a high *a priori* potential for sustained host colonization, and then requires empirical testing to confirm that such persistent candidates deliver a measurable host benefit. The proof-of-concept lies in demonstrating that a candidate (MC10-B4) chosen specifically for its genomic signatures can achieve sustained retention and confer beneficial host effects in a real-world setting. Consequently, the field trial was designed to include a placebo control. The objective was not to benchmark MC10 against alternative candidates, but to test whether an evolutionarily informed selection framework can prospectively identify a probiotic with a high probability of sustained colonization and functional benefit under field conditions. A successful outcome validates the framework as a strategy for de-risking probiotic selection.

We performed a controlled laboratory experiment to rule out alternative explanations for a potential probiotic benefit. The nutritional contribution of the inoculum was negligible because the total dose delivered (∼10^7^ cells/mL per fragment twice a week over four weeks) was orders of magnitude lower than the ambient bacterial load in the rearing system (∼3 × 10^11^ cells in a 200 L volume at ∼1.5 × 10^6^ cells/mL). MC10-B4 established at significantly higher levels in *Acropora pruinosa* than the sympatric control strain MC15-BG7, confirming that this colonization capacity was aligned with MC10’s genomic predisposition for persistent residency, rather than being a general trait of coral-associated *Ruegeria*. Live inoculation with either MC10-B4 or MC15-BG7 did not alter baseline coral physiology across physiological metrics, including dark-adapted maximum photosynthetic efficiency, coloration, host protein content, total lipid energy content, chlorophyll a concentration, symbiont density, and host fluorescent protein content, compared to an AFSW placebo, which ruled out an inoculation-induced priming effect, independent of sustained colonization, as the cause of later field-observed benefits.

The field trial tested the core prediction that persistent colonization by MC10-B4 would enhance host thermal resilience. Treatments were deployed at separate but proximate sites (∼40 m apart) within the same habitat, minimizing the risk of cross-contamination via passive dispersal while ensuring similar environmental conditions (depth 4 m; sand-rubble substrate; tide-driven flow, 0.2±0.3 m/s, (*31*). At this scale, water depth, rather than along-reef distance, primarily structures environmental variability on coral reefs (*32–34*). Baseline monitoring confirmed that key abiotic parameters, including water temperature and dissolved inorganic nitrogen, were homogeneous between sites (**Fig. S9**). The primary environmental driver during the observation period was an extreme, regional heat stress event (DHW 8.8-18 °C-weeks).

To exclude postbiotic effects, we compared coral responses to live, autoclaved, and pasteurized bacterial cells. This experiment revealed two distinct outcomes. Exposure to pasteurized MC10-B4 cells, which are expected to preserve some heat-labile metabolites, reduced the maximum photosynthetic efficiency. This effect contrasts with the relative preservation of photosynthetic efficiency observed in MC10-inoculated corals under field stress. This result indicates that a one-time release of heat-labile metabolites cannot explain the beneficial outcome observed in the field. The same treatment increased host fluorescent protein content, a response associated with antioxidant capacity. One interpretation is that specific MC10 metabolites could initiate a beneficial stress-response priming in the holobiont. These responses were not observed with live MC10-B4, nor with pasteurized MC15-BG7 or autoclave-killed cells. Therefore, even if certain metabolites could theoretically prime the host, for such priming to provide protection during a bleaching event months later would require the continual presence of the metabolite source: a persistently resident bacterial population. Thus, any integrated ‘probiotic-postbiotic’ mechanism fundamentally depends on the sustained colonization that our evolutionary framework selects for.

Collectively, these experiments argued against mechanisms relying on nutrition, transient priming, or one-time postbiotic effects. The treatment-specific benefits observed during peak thermal stress, supported by mechanistic dissection, indicate that sustained colonization was the underlying cause. This validates the core prediction of our evolution-guided framework.

### Facultative endosymbionts as host-generalist probiotics

*Ruegeria* MC10 can be described as a facultative endosymbiont, given its variable occurrence among conspecific coral colonies and its detection in free-living environments. Such facultative symbionts exhibit a distinctive duality: genomic erosion drives metabolic dependency whereas ecological flexibility is retained. This life history strategy may enable a broad host range.

MC10’s host versatility could arise from mechanisms that exploit conserved host interfaces rather than lineage-specific adaptations. For instance, strain MC10-B4’s pseudogenized TCA cycle necessitates scavenging host-derived metabolites like succinate or fumarate, pathways conserved across eukaryotes. This strategy aligns with “resource tracking”, a mechanism whereby symbionts exploit universal host resources to colonize diverse hosts (*35*). Colonization could also involve interactions with conserved eukaryotic cell interfaces, a concept reminiscent of generalist probiotics like *Lactiplantibacillus plantarum*, which uses a conserved colonization island to inhabit hosts from *Drosophila* to humans (*36*). This aligns with “ecological fitting”, where pre-existing traits enable colonization of novel hosts without coevolution, a key driver of host generalization in facultative symbioses (*35*).

MC10’s persistence in the environment alongside corals could reflect relatively weak host selection pressures. When hosts impose minimal filtering, symbionts face reduced costs of adaptation to novel hosts, facilitating niche expansion (*35*). MC10’s low relative abundance in coral microbiomes likely minimizes its burden on the host, potentially weakening selection for specialization. The scalability of this probiotic approach is enhanced by such generalism. Unlike obligate symbionts restricted to single hosts (*10*), facultative endosymbionts like MC10 may exploit conserved molecular gateways, enabling nursery-based priming of diverse coral species with a single probiotic preparation.

### Microbiome dynamics and stability

A central goal of probiotic intervention aligns with microbial transfer therapy in human medical settings: introducing beneficial taxa to reset dysbiotic microbiomes and enhance resilience through targeted restructuring (*20, 37*). Using *Ruegeria* MC10 as a keystone inoculant, we observed microbiome modulation without long-term destabilization. While β-diversity differences between probiotic-inoculated and placebo-treated corals peaked post-inoculation, communities converged over the monitoring period, indicating environmental acclimatization.

During the peak of bleaching stress (**t6**), the presence of MC10-B4 was associated with a specific restructuring of the coral microbiome. Analysis revealed significant enrichment of Bacteroidetes taxa specializing in polysaccharide degradation (*Cryomorpha*, *Spirosoma*, *Sunxiuqinia*, *Gelatiniphilus*, *Aureispira*, *Flavobacterium*) (*38*), ROS-tolerant genera (*Deinococcus*, *Desertibacter*, *Spirosoma*) (*39*), the *Vibrio*-predatory specialist (*Aureispira*) (*40*), and known probiotic taxa (*Veillonella*, *Limosilactobacillus*) (*41, 42*). Concurrently, multiple Roseobacteraceae genera were depleted (*Sulfitobacter*, *Roseovarius*, *Pseudoruegeria*, *Donghicola*, *Aliiroseovarius*, *Phaeobacter*). This depletion may indicate competitive restructuring within this family or may reflect their close association with Symbiodinaceae (*43*), which are themselves stressed and declining during bleaching. These shifts were largely absent before (**t3**) and after (**t8**) the bleaching event, indicating that the earlier MC10 inoculation (at **t0**) was associated with the establishment of a microbiome state potentially more favorable for stress response.

Algal symbiont switching (e.g., from sensitive *Cladocopium* to resilient *Durusdinium*) represents a known thermal adaptation pathway in some corals (*44*), but this mechanism is unlikely to explain our results. Region-wide characterization confirms that Hong Kong corals, including *Acropora* spp., predominantly harbor *Cladocopium* spp. (except *O. crispata*) (*45*). This consistent symbiont profile across host species and sites makes algal-mediated thermal tolerance an improbable alternative explanation for the enhanced resilience observed.

### Towards scalable probiotic interventions

Coral restoration commonly involves growing fragments in nurseries before outplanting (*46, 47*). This operational framework is highly compatible with integrating probiotic intervention as a nursery-applied treatment. Our study establishes a pipeline where evolutionary genomics first selects candidates with a high *a priori* potential for long-term host residency. We employed CBASS as a rapid, reproducible assay to obtain an initial functional readout of acute thermal tolerance, as reported in our companion study (*21*). This step helps triage candidates before resource-intensive field trials.

Our 8-month field trial provides a proof-of-concept that evolutionarily selected probiotics can achieve sustained retention and be associated with enhanced stress tolerance during a natural bleaching event. Critical knowledge gaps remain for evaluating long-term restoration success, an issue plaguing coral restoration initiatives globally (*47*). Multi-year data on coral survival, reproductive fitness, and reef-scale outcomes such as biodiversity recovery and structural complexity are essential next steps.

The scalable application of any probiotic requires a favorable ecological risk profile. *Ruegeria* MC10 occurs naturally across multiple coral species, seawater, and sediment, indicating ecological flexibility without rampant proliferation. Several convergent factors suggest a low-risk, self-limiting phenotype aligned with its selected genomic trajectory: consistently low natural abundance, a pseudogenized metabolism that constrains competitive growth outside host tissues, and minimal field-established abundance within coral gastrodermis. The microbiome restructuring observed during bleaching was a transient, fine-scale modulation rather than a dysbiotic takeover. This combination of traits defines a probiotic candidate that integrates into the holobiont without disruptive dominance, a key consideration for safe, multi-species restoration.

The beneficial effects of MC10-B4 in both a model cnidarian (*21*) and a reef-building coral suggest potential for cross-species efficacy. Further validation across a wider range of coral species, coupled with rigorous evaluation of potential off-target effects, is a necessary next step. Developing high-efficiency, scalable delivery technologies for underwater nursery applications could improve treatment effectiveness. If this approach proves generalizable and safe, such broad-host applicability would significantly enhance scalability. This would enable the same probiotic preparation to be used across multiple coral species within nursery-based workflows. This initial proof-of-concept points toward a promising avenue for developing scalable, microbial-based interventions to help bridge the resilience gap for coral reefs.

## Acknowledgements

We thank all the colleagues who have supported experiments to test physiological priming effect, outplant fieldwork, and FISH experiments, including Jingqi Gao, Kai Yan Lai, and Nikolai B. Tang. We would like to thank Kristin Bär-Häge and Myriam Schmid for their support with Aiptasia animal husbandry for CBASS acute thermal assays.

## Author Contributions

Conceptualization: H.L. Methodology on *Ruegeria* isolation, genome analysis, and population-level primer design: M.X., T.L., H.L. Methodology and investigation on CBASS: N.X., M.D., H.M., C.R.V. Code development: T.L., X.W. Investigation on coral sampling and *Ruegeria* isolation: M.X., K.E.H., D.L., S.E.M., R.H., E.C., Z.W., P.T., J.B. Investigation on bacterial growth rate: C.J.X., Q.H. Methodology and investigation on probiotic priming and postbiotic effect: N.X., P.L., C.T.C., M.X., H.L., R.S.P., C.R.V., A.P.Y.C. Field trial: A.P.Y.C., C.T.C., C.H.L, W.Y.T, K.K.T. Methodology on FISH: Z.X., G.C. Data visualization and statistical analysis: N.X., S.E.M. Funding acquisition: H.L., A.P.Y.C., C.R.V. Project administration: H.L. Supervision: H.L., C.R.V., A.P.Y.C. Writing – original draft: H.L. Writing – review and editing: C.R.V., H.L., N.X., M.X., S.E.M., R.S.P., G.C. Writing – supplementary materials: M.X., N.X., T.L., C.T.C., A.P.Y.C., H.L.

## Data and materials availability

All raw sequencing data generated in this study have been deposited in the National Center for Biotechnology Information (NCBI) database. Whole-genome next-generation sequencing data for *Ruegeria* isolates, including raw reads (n = 409) and genome assemblies (n = 419), are deposited under BioProject accession PRJNA1264799. Nanopore sequencing raw reads and assemblies for 34 *Ruegeria* isolates are available under BioProject accession PRJNA1275854. Metabarcoding *ATP5B* and *parC* amplicon sequences for natural coral samples are available under BioProject accessions PRJNA1275576 and PRJNA1275610. Metabarcoding 16S rRNA and *parC* amplicon sequences for outplanted corals are available under BioProject accession PRJNA1275585 (https://dataview.ncbi.nlm.nih.gov/object/PRJNA1275585?reviewer=9ne88342f7at8ndur1klq05 mo8) and PRJNA1275617 (https://dataview.ncbi.nlm.nih.gov/object/PRJNA1275617?reviewer=e6g3aatk7pg8r6m6ao0riovj nu), respectively. All data will be publicly accessible upon publication. The scripts used for genome assembly and annotations, population genomic analysis, amplicon data analysis, and pseudogene analysis have been deposited in an online repository (https://github.com/444thLiao/CoralProbiotic).

