## Supplementary information for "Evolutionary genomics predicts probiotic persistence in corals"

#### **The PDF file includes:**

#### Materials and Methods

#### Figs. S1 to S9

#### **Other Supplementary Materials for this manuscript include the following:**

#### Tables S1 to S6

#### Datasets S1 to S2

### Materials and Methods

#### 1. *Ruegeria* isolation and population genomic analysis

##### 1.1 Coral sampling, bacterial isolation, and genome sequencing

Five coral species were sampled from multiple Hong Kong reef sites for *Ruegeria* isolation (Fig. S1**).** *Platygyra acuta* and *Acropora digitifera* were sampled from Kiu Tsui Chau, Wong Wan Chau, Chek Chau, and Ngo Mei Chau between April 2017 and February 2019. *Oulastrea crispata* was sampled from Bluff Island, Lo Chau, Sham Wan, and Yam Tsai Wan in December 2020. *Acropora* *solitaryensis* and *Favites pentagona* were sampled from Ninepin Islands in June 2021. Two coral mother colonies (~5 cm²) were collected for each species from each site at a depth of 2-3.5 meters using a hammer and chisel.

On the sampling day, three coral compartments (mucus, tissue, and skeleton) were separated for *Ruegeria* isolation. Coral fragments were washed three times with autoclaved 0.22 μm-filtered seawater (AFSW) to remove loosely attached microbes on coral surfaces. Mucus was collected using a 100 μL pipette, tissue was flushed using a Waterpik (Waterpik, US) with AFSW, and skeleton was pulverized in grain size and resuspended in 1 mL of AFSW. Mucus, tissue, and skeleton slurries were serially diluted (10-fold) and spread on marine basal medium (MBM) agar plates. The medium recipe was modified from a previous protocol (*1*) by replacing the nitrogen source with glycine betaine and N-acetylglucosamine to achieve a higher *Ruegeria* isolation efficiency. Plates were incubated at 28 °C for two weeks, and single colonies were picked and purified three times on Marine Agar 2216 (BD Difco, USA). Taxonomic identification was performed using full-length 16S rRNA gene sequencing, with reference to EzBioCloud database (*2*).

Genomic DNA from 419 *Ruegeria* isolates was extracted using the TIANamp Genomic DNA Kit (TIANGEN Biotech, China) following the manufacturer’s instructions. DNA libraries were constructed using the NEBNext® Ultra™ II FS DNA Library Prep Kit (E6177S, NEB, US) and subject to next-generation sequencing (NGS) on the DNBseq PE150 platform (BGI, China). Subsequent data processing was carried out as previously described (*3*). Raw reads were trimmed using Trimmomatic (*4*) and reads shorter than 50 bp were discarded. Quality assessment was performed using FastQC (*5*), and clean reads were assembled into contigs using SPAdes (*6*).

##### 1.2 Phylogenomic and population genomic analysis.

A total of 445 genomes were analyzed, comprising draft genomes from 419 newly isolated *Ruegeria* strains and 26 reference genomes from NCBI GenBank, for phylogenomic and population genomic analysis (Dataset S1). Orthologous gene families were identified using OrthoFinder v2.5.1 (*7*), and single-copy orthologs were aligned using MAFFT v7.471 with the “auto” mode (*8*). Alignments were trimmed using trimAl v1.4 with the “automated1” mode 4 (*9*). A maximum-likelihood phylogenomic tree was constructed using IQ-TREE v1.6.12 with the best-fit model (JTT+I+G4) selected by ModelFinder (*10*). Node support was assessed using 1,000 ultrafast bootstrap replicates. All trees were edited and displayed with iTOL v5 (*11*). Pairwise 16S rRNA gene identity and genome-wide average nucleotide identity (ANI) were calculated using BLAST 2.6.0+ (*12*) and fastANI (*13*), respectively.

Populations were delineated using PopCOGenT (*14*), which identifies evolutionarily cohesive clusters (main clusters, MCs) by analyzing genome-wide allele sharing across isolates’ or single-cell genomes. Unlike methods dependent on fixed thresholds (e.g., average nucleotide identity or ANI 95%) or tools building on the ratio of homoplastic and non-homoplastic alleles, which fail to distinguish recent gene flow from historical events, PopCOGenT solves this issue while aligning with the biological species concept. In *Prochlorococcus*, MCs were validated as monophyletic groups in most single-copy gene trees, confirming genetic cohesion (*15*). PopCOGenT defines populations by gene flow barriers, contrasting ecological classifications based on habitat co-occurrence rather than reproductive isolation.

##### 1.3 Nanopore sequencing, genome assembly, gene prediction and annotation

Long-read Nanopore sequencing was performed for 34 representative isolates from 19 MCs to generate closed genomes that serve as reference sequences for each identified MC. These 19 MCs are phylogenetically diverse, and each has at least three isolates, except for MC46 which has two isolates. MC10 has 15 closed genomes because it exhibited a uniquely high number of contigs following NGS. MC46, i.e., the sister group of MC10, has two closed genomes. There were 17 closed genomes for the remaining 17 MCs each. DNA libraries were prepared using a Ligation Sequencing Kit (SQK-LSK110, Oxford Nanopore Technologies) and sequenced on a MinION sequencer with an R9.4.1 flow cell. Long-read assemblies were performed using i) the hybrid mode of Unicycler (v0.5.0) using an overlap-layout-consensus (OLC) - based method (*16*), ii) Flye (v2.6) (de Bruijn graph) (*17*), and iii) Canu (v2.2) (OLC-based method) (*18*). Assemblies were polished using Pilon v1.24 (*19*) based on NGS short reads (Dataset S2). Assembly quality was evaluated using Mauve (*20*) and Bandage (*21*), and circularized genomes were adjusted using Circlator (*22*) to place the *dnaA* gene at the start.

Protein-coding genes were predicted using Prokka v1.14.6 (*23*). Genome completeness was assessed using CheckM v1.0.7 (*24*) and miComplete v1.1.1 (*25*). Functional annotation was performed using KOfamscan (*26*), COG (*27*), and TIGRFAM databases with HMMER v3.3 (with e-value cutoff of 1e-20) (*22*).

#### 2. Analysis of insertion sequences, pseudogenes and *Ruegeria* growth rate

##### 2.1 IS identification and analysis

Insertion sequences (ISs) were predicted in 34 newly closed *Ruegeria* genomes and the reference genome *Ruegeria* *pomeroyi* DSS-3 downloaded from NCBI GenBank database using Insertion sequences (ISs) were identified using ISEscan v1.5.4 4 (*28*). IS families were classified as described in previous study (*29*).

##### 2.2 Pseudogene identification, classification and functional prediction

Pseudogenes were predicted using Pseudofinder (*30*) against the NCBI Non-Redundant protein database (accessed on Sept. 29, 2022) (*31*). Pseudogenes shorter than 300 nt were excluded (*32*). Pseudogenes overlapping single ORFs were assigned to the function of the overlapping ORF.

Pseudogenes were classified into four categories, “IS disruption”, “Frameshift mutations”, “Nonsense mutations”, and “Singletons”, to understand the evolutionary processes behind pseudogenization. Pseudogenes with both mutation types were tagged as “Unclassified”. The classification was based on pairwise BLASTP alignments across the genomes analyzed, using default parameters. Pseudogenes overlapping ISs and aligned to intact genes in other genomes were designated as “IS disruption”. In instances where no IS was identified, pseudogenes were categorized based on their genetic variation relative to the aligned intact gene. If only point mutations were present, these pseudogenes were classified as “Nonsense mutations”. In contrast, if only frameshift mutations were detected, they were classified as “Frameshift mutations”. Finally, pseudogenes that could not be aligned with any intact gene were classified as “Singletons”.

##### 2.3 Evaluation of metabolic pathway completeness and pseudogenization of associated genes

Metabolic pathway completeness was evaluated using KEGG ortholog profiles and KOfamscan (*26*) along with the metabolic completeness definitions implemented in MicrobeAnnotator (*33*). Genes not found by KOfamscan were re-annotated using InterProScan (*34*). If all genes annotated to a specific KO category were intact based on KOfamscan or InterProScan, the KO was considered present in the *Ruegeria* genome. If all associated genes were pseudogenes, each pseudogene was manually assessed to confirm truncation status using DNA Feature Viewer (*35*).

##### 2.4 *Ruegeria* growth kinetics analysis

Growth kinetics of 24 *Ruegeria* isolates were compared. The set included three strains from the population MC10, two from its sister population MC46, 18 strains from other *Ruegeria* MCs, and the free-living model strain *R. pomeroyi* DSS-3. All strains were cultured in Marine Broth 2216 at 28 °C with shaking at 200 rpm. Growth was monitored by measuring the optical density at 600 nm (OD600) over 72 h. Biological triplicates were performed for each strain. The maximum growth rate (μ_max_) for each strain was calculated based on the slope of natural log-transformed OD_600_ values during the exponential growth phase (i.e., from 4 h to 8 h). Additionally, the maximum biomass yield of each strain was evaluated through colony-forming unit (CFU) counts at the stationary phase (i.e., at 72 h). To account for the phylogenetic non-independence of *Ruegeria* isolates and confounding effects of their evolutionary origins on growth rate, we performed the phylogenetic analysis of variance (PhylANOVA) using the “phylANOVA” function in the R package “phytools” (v2.4). The model included one fixed factor with “if MC10” as the primary predictor and MC identity as a covariate. The analysis was conducted with 10,000 simulations to generate a null distribution of the F-statistics under the Brownian motion evolution model. A phylogenomic tree for 24 *Ruegeria* isolates used in the growth kinetics analysis was implemented in the PhylANOVA analysis.

#### 3. Preliminary survey of MC10 in natural corals and free-living niches in Hong Kong reefs

Amplicon sequencing of the two marker genes (*ATB5B* and *parC*) was performed to survey the relative abundance of MC10 in natural corals and surrounding environments. 44 coral fragments of five coral species (*Oulastrea crispata*, *Acropora solitaryensis*, *Plesiastrea versipora*, *Porites* sp., and *Platygyra acuta*) were sampled from five Hong Kong reef sites and stored at -80 °C until processing. For free-living niches, 1 L of seawater and/or 5 g of sediment were collected from three reefs at Crescent Island, Sharp Island, and Wu Pai. Seawater was filtered through a 0.22 μm membrane and the membrane was stored at -80 °C until processing. DNA extraction was performed using the DNeasy PowerSoil Pro Kit (Qiagen) and amplicon sequencing of two genes was conducted on the DNBseq PE300 platform (BGI, China). Amplicon data was analyzed to determine the relative abundance of MC10 following the methodology described previously (*36*).

#### 4. An 8-month field trial with *Ruegeria* MC10 in Hong Kong

##### 4.1 Comparative analysis of colonization efficiency between MC10 and MC15

Prior to field outplanting, we evaluated the colonization capacity of next-generation probiotic MC10 versus a conventional probiotic, *Ruegeria* population MC15. Strains MC10-B4 and MC15- BG7 were used to control for ecological background of the strains because they were isolated from the same coral colony. Bacterial inocula were prepared by culturing single colonies in Marine Broth 2216 (BD Difco, USA) at 28°C and 280 rpm to reach an OD_600_ of 0.8. Bacterial suspensions were centrifuged at 3,000 g for 5 min, washed three times with AFSW, and diluted 100-fold to a final concentration of 10^7^ CFU/mL.

A total of 15 coral fragments (5 colonies × 3 replicates) of *A. pruinosa* were inoculated with either placebo AFSW, MC10-B4, or MC15-BG7 twice weekly for one month. Before inoculation, fragments were exposed to air for 2 min to facilitate mucus release. Each fragment received 1 mL of inoculum using mucus as a delivery vector and was incubated for 2 min before being returned to the main tank (*37*). To minimize the effects of water flow, circulation pumps were turned off for 4 hours after the inoculation.

Post-inoculation for one month (**t0**), all 45 fragments were collected and stored at -80 °C for downstream analysis. Coral tissue was washed off using a WaterPik with 50 mL of AFSW, concentrated by centrifugation at 3,000 g for 10 min at 4°C, and processed for DNA extraction with the DNeasy PowerSoil Pro Kits (Qiagen, Germany). Extracted DNA was quantified using a Qubit fluorometer and the Qubit^TM^ dsDNA HS Assay Kit (Invitrogen, USA).

To assess the relative abundance of MC10 and MC15 in the *Ruegeria* community and in the total bacterial community of coral holobionts, we employed a dual-marker amplicon sequencing approach combining 16S rRNA gene amplicon sequencing (V5-V6 region) and *Ruegeria* population-resolving *parC* gene amplicon sequencing. For 16S rRNA gene amplicon sequencing, the primer set 784F (5’-TCGTCGGCAGCGTCAGATGTGTATAAGAGACAGAGGATTAGATACCCTGGTA-3’) and 1061R (5’-GTCTCGTGGGCTCGGAGATGTGTATAAGAGACAGCRRCACGAGCTGACGAC-3’) was used (*38-40*). For *parC* gene amplicon sequencing, the primer set 668F and 1189R was used (Table S2). The amplicon data processing workflow followed the methods described previously (*36*).

Statistical analysis for relative MC10 and MC15 abundances in *Ruegeria* and total bacterial communities was conducted using the linear mixed models (LMMs) implemented in the lme4 package in R version 4.1.1 (*41*). This model included one fixed effect (treatment) and one random effect (colony) to account for the individual variability. We graphically checked for all models whether the residuals were normally distributed using the histograms and quantile-quantile plots. The homoscedasticity of the residuals was checked by plotting them against the fitted values and each explanatory variable. The significance of the fixed effect was evaluated via one-way ANOVA.

##### 4.2 One-month coral inoculation with live and heat-killed *Ruegeria* MC10

To determine whether the observed beneficial effects of *Ruegeria* MC10-B4 in field trials were mediated by mechanisms independent of sustained colonization, we conducted a controlled, one-month inoculation experiment. This assay was designed to differentiate between two non-colonizing modes of action: (1) transient, broader microbiota-priming effects, which could include nutritional supplementation or other community-level modulation, and (2) postbiotic effects mediated by microbial cellular components and metabolites from inactivated cells. Using a crossed factorial design, we compared the strain of interest, MC10-B4, to a sympatric control strain, MC15-BG7, which was isolated from the same coral host but lacks the genomic signatures of host dependency.

The experiment comprised seven treatments: placebo control, live MC10-B4, pasteurized MC10-B4, autoclaved MC10-B4, live MC15-BG7, pasteurized MC15-BG7, and autoclaved MC15-BG7. For this, five mother colonies (i.e., genotypes) of *Acropora pruinosa* were collected from Bluff Island (22°19'31.9"N 114°21'14.9"E) in Hong Kong SAR. After a 3-day acclimatization in the 200 L main tank (temperature: 23.07 ℃ ± 0.50 ℃, salinity: 31 ± 0.50 ppt, light: ~60 μmol m^-2^ s^-1^, water flow: ~20 L h^-1^), corals were cut to a uniform size (~5 cm in length) and left to recover for 5 days before bacterial inoculation. In total, 105 coral fragments (5 mother colonies × 7 treatments × 3 biological replicates) were kept in 35 (3 fragments from the same mother colony per tank) of individual 4 L aquarium tanks. These aquarium tanks were supplied with continual air and fresh seawater that was pre-filtered via a 50 µm water filter cartridge to exclude the plankton. The flow rate of 0.76 L/h in the 4 L tank led to a daily water exchange of ~18.24 L of fresh seawater.

Single colonies of *Ruegeria* strains MC10-B4 and MC15-BG7 were picked and cultivated in Marine 2216 Broth (Difco, USA) for 48 hours until the late-log phase (~10^9^ CFU mL^-1^). Bacterial cells were collected, washed three times with the autoclaved 0.22 μm-filtered seawater (AFSW) (3,000 g, 5 min), and resuspended at a final concentration of 10^7^ CFU mL^-1^. Apart from one aliquot for live bacterial cells, another two aliquots were killed using either pasteurization (70 °C, 30 minutes) (*42*) or autoclaving (121 °C, 20 minutes) (*43*), respectively. The inactivation of *Ruegeria* cells was confirmed by no visible colony on the Marine 2216 Agar. It is notable that, compared to the autoclave method, pasteurization represents a less extreme treatment limiting the denaturation of bacterial cellular components (*42*). To inoculate the coral fragment, 1 mL of *Ruegeria* suspension (or AFSW for placebo) was applied directly onto the coral surface area, using the mucus as a delivery vector. Coral fragments were exposed to air for 3 minutes to induce the mucus release, then inoculated with the target *Ruegeria* suspension (or AFSW for placebo), and held for 2 minutes before being returned to the tanks. To optimize the host-microbial interactions post-inoculation, water flow and aeration were stopped for 6 hours after the inoculation and then resumed to normal conditions. Any additional feeding was abandoned throughout the experiment to avoid confounding effects from additional nutrient uptake via heterotrophy.

Physiological metrics were measured for corals at the end of the nutritional experiment (i.e., on day 27). Maximum photosynthetic efficiency (F_v_/F_m_) was measured for corals after 30 minutes of dark acclimation. Coral coloration is determined with the Coral Health Chart (*44*). Coral tissue was sprayed off from the skeleton for each fragment using a high-pressure airbrush (RoHS, China) with pre-cold AFSW. Coral tissue was homogenized at 3500 rpm for three times (on ice, 30 s intervals in-between) with T-18 UltraTurrax (IKA, Germany). Coral homogenates were used for centrifugation at 3000 g, 5 min, at 4 °C. The supernatants were collected for host protein determination and measurements of the relative fluorescence unit. Host protein was measured with the BCA Protein Assay Kit (Tiangen, China) according to the instructions. The Relative fluorescence unit was determined following a previous study (*45*), with the emission spectra of each sample measured using 384-well black/clear microtiter plates with 20 μl of sample/well in triplicate. Each well was excited at 280 nm, and the emission was recorded between 460 and 600 nm. The total fluorescence was calculated between the emission spectrum 465 and 600 nm for each sample. Symbiont cells were washed in three cycles of centrifugation (3000 g, 5 min) and resuspension in 1 ml of AFSW. For symbiont density, two aliquots of 10 μl Symbiodiniaceae homogenates were used for Symbiont density counting using the hemocytometer slide and microscope (Olympus CKX53, Japan). For chlorophyll a content, 1 ml of washed symbiont cell suspension was transferred into an Eppendorf tube, pelleted, and resuspended in acetone (100%). Samples were incubated in the dark at 4 °C for 24 h before three technical replicates of 200 µL were transferred into a 96-well plate. The absorption of samples was immediately recorded at 630, 664, and 750 nm using the BioTek Epoch 2 Microplate Spectrophotometer (Agilent, USA). Chlorophyll a content was calculated following the study of (Jeffrey and Humphrey). Coral surface area was determined with the “aluminum foil” method (*46*). Host protein, algal symbiont density, and chlorophyll a content was corrected for sample volume and normalized to the coral surface area. Total lipid extraction of coral holobionts and subsequent energy content analysis of the extracted lipid (i.e., total lipid energy content) were conducted according to a published protocol (*47*). Snap-frozen coral fragments were ground to powder with the liquid nitrogen, freeze-dried overnight with a freeze dryer (Labconco, USA), and stored at -80 °C until further analysis. Extraction solvent (chloroform:methanol, 2:1 v/v) containing 50 mg/L butylated hydroxytoluene (BHT) was prepared fresh daily. Approximately 0.5 g of ground coral holobiont powders per sample was extracted with 10 mL extraction solvent in a glass vial. Skeletons were removed by vacuum filtration (GF/F filters), with filtrates washed with 0.88% KCl in a separatory funnel. The organic phase was collected from the funnel, air dried with N_2_ (100% purity, Linde), and weighed with an analytical balance (A&D company, Japan), which is the lipid weight. Residual skeleton for each coral sample was weighed before and after burning at 450 °C for 5 h in a muffle furnace (Lindberg, USA), the difference between which referring to ash-free dry weight (AFDW). The energy content of lipid extracted (i.e., total lipid energy content) was calculated from lipid weight (*47*) and normalized to AFDW (*48*). Visualization and statistical analysis for coral physiological data were conducted in R v4.1.0 with several packages, such as “ggplot2”, “lme4”, and “tidyverse” (*49-51*). We used linear mixed models (LMMs) to test the statistical significance across samples by defining one fixed effect (treatment) and two random effects (colony identity and tank ID). The significance of the fixed effect was evaluated via a one-way ANOVA and Tukey’s HSD test.

##### 4.3 Coral outplant and physiological monitoring

The sensitive branching coral *A. pruinosa* was selected as the target species for the field trial due to its ecological significance in reef-building, global prevalence, and dramatic decline in many reefs, including western Hong Kong reefs exposed to excess nutrient loading (*52, 53*). Five mother colonies of *A. pruinosa* were sampled from Bluff Island (Hong Kong) and transferred to the Marine Science Laboratory at CUHK. Corals were reared in 300 L aquarium tanks with flow-through seawater pre-filtered through a 50 μm mesh bag. Temperature and light conditions were continuously monitored using a HOBO data logger (Onset Computer Corporation, USA). Water temperature was maintained at 23 ± 3 °C, and natural light was supplemented with shading. Coral mother colonies were fragmented into 4-5 cm pieces and acclimated for two weeks before *Ruegeria* inoculation. The coral inoculation procedure followed the standardized protocol detailed in Section 4.1. Given the superior colonization efficiency in corals, MC10-B4 was selected for the field trial to test its persistence and putative probiotic efficacy in corals. In contrast, MC15-BG7 was excluded due to its poor colonization in corals after one-month inoculation.

Prior to outplanting, coral fragments were secured onto customized platforms (~50 cm width × 55 cm length × 30 cm height), which were constructed from PVC pipes and designed with checkerboard coral plugs (Fig. S6). Two platforms were utilized, each with four checkboards, including one for 60 fragments of placebo-treated group and the other for 60 fragments of *Ruegeria* MC10-B4-inoculated group. The platforms were deployed at a depth of ~ 4 meters, with 40-meter spacing to minimize cross-contamination between treatments. The platforms loaded with coral fragments were deployed at Knob Reef (22°27’48.9” N, 114°17’28.3” E), where they remained for an 8-month (January-September 2024) monitoring period to assess probiotic performance under natural reef conditions. Critically, the 2024 summer thermal event constituted a mass regional bleaching episode (DHW = 8.8-18 °C-weeks), with a vast majority of shallow-water corals across Hong Kong reefs exhibiting visual bleaching. This uniform regional stressor ensured both treatment platforms experienced near-identical thermal regimes, despite their 40-meter spacing.

Coral health metrics (i.e., size, coloration, and *in situ* operational efficiency of PSII under ambient light conditions F_v_′/F_m_′) were assessed at 1 (**t1**), 3 (**t3**), 6 (**t6**), 7 (**t7**), and 8 (**t8**) months post-outplanting. High-resolution photos were taken underwater for corals using the digital camera (Tough TG-6, Olympus). Coral coloration was determined for each fragment under consistent lighting conditions using the CoralWatch Coral Health Chart following the protocol of (*44*). Concurrently, F_v_′/F_m_′ was measured for each coral fragment underwater using the Diving-PAM (Walz, Germany). Despite that coral fitness metrics mentioned above were consistently measured at 3 p.m. on each sampling day, F_v_′/F_m_′ is not appropriate to be compared across timepoints as it is affected by natural light variances. For molecular analysis, 15 fragments (5 mother colonies × 3 replicates) from the MC10-B4-inoculated and placebo-treated groups were collected after 1, 3, 6, and 8 months post-outplanting. Given the destructive nature of molecular sampling, which precludes repeated measures on the same coral fragments, invasive analyses were restricted to four critical phases: field acclimation (**t1**), pre-bleaching (**t3**), during bleaching (**t6**), and post-bleaching recovery (**t8**). This conserved coral biomass while enabling focused assessment of MC10’s probiotic effects across thermal stress transitions. Phenotypic metrics (e.g., coloration, photosynthetic efficiency) were collected at all intervals, including **t7**, but molecular analyses (e.g., microbiome profiling) were confined to the selected timepoints. Collected samples were transported to the Marine Science Laboratory of CUHK and stored at -80 °C until further analysis.

All statistical analyses were conducted in R version 4.1.1 (*41*) packages “ggplot2” (*54*) and “lme4” (*55*). For coral coloration, we used linear mixed models (LMMs) defining three fixed effects (*Ruegeria* treatment, time, and their interactions) and one random effect (colony and time interactions). For F_v_′/F_m_′, we used LMMs defining two fixed effects (*Ruegeria* treatment, interactions of time and *Ruegeria* treatment) and one random effect (colony and time interactions). Significance of fixed effects was evaluated via two-way ANOVA and Tukey’s HSD. We graphically checked for all models whether the residuals were normally distributed using the histograms and quantile-quantile plots. The homoscedasticity of the residuals was checked by plotting them against the fitted values and each explanatory variable.

##### 4.4 16S rRNA gene amplicon sequencing and differentially abundant taxa in MC10-B4-treated corals

Bacterial community structures between placebo and MC10-B4-treated corals across five timepoints: baseline (**t0**) and 1, 3, 6, and 8 months post-outplanting (**t1-t8**) was visualized using the Principal Coordinates Analysis (PCoA) and Bray-Curtis distances. The similarity of bacterial community structures between treatments across timepoints was assessed via ANOSIM and Bray-Curtis distances.

Analysis of Compositions of Microbiomes with Bias Correction (ANCOM-BC) was used to identify differentially abundant taxa at the genus level between placebo and MC10-B4-treated corals at **t6** (i.e., during bleaching). ANCOM-BC (v2.1+) was conducted based on an ASV table aggregated to the genus level (tax_level = “Genus”). After filtering to retain taxa present in ≥5% of samples (prv_cut = 0.05), we tested the treatment effects while controlling for false discovery rate (Benjamini-Hochberg method, p_adj_method = “BH”) and accounting for compositional biases (lib_cut = 1000; struc_zero = TRUE for zero-inflation correction). Differential taxa abundance between placebo and MC10-B4-treated corals was assessed through log2-fold changes (log2FC), with significance defined as FDR-adjusted p-values < 0.05. For visualization, we generated a heatmap displaying log2-transformed read counts (log2[reads+1]) of the top 20 significantly upregulated and downregulated genera (ranked by absolute log2FC magnitude), respectively. The pseudocounts were added to handle the zero case in the calculation.

##### 4.5 Fluorescence in situ hybridization (FISH) of *Ruegeria* pure cultures with a *Ruegeria* MC10-specific probe

To enable precise discrimination of *Ruegeria* MC10 population from other populations using fluorescence in situ hybridization (FISH), we developed a highly specific probe targeting a 215-bp region of the population-resolving *parC* gene that is uniquely conserved in MC10. The probe was designed against the following amplified sequence:5’- TCCGCGACAACTATGACGGCACGCTGACCGAGCCGGTCGTGCTGCCGGCGCAGTTCCCCAACCTGCTGGCCAACGGTGCCAGCGGTATCGCGGTGGGCATGGCGACCAACATTCCGCCGCATAACATCTCGGAGCTCTGCGATGCCTGTCTGCACCTGATCAAAACGCCGGACGCGCGTGACGACACGCTATTGAACTACGTTCCCGGTCCTGAC-3’.

For validation of FISH probe specificity, pure cultures of MC10-B4 and MC15-BG7 were prepared. MC15-BG7 served as a control for the MC10-specific probe. Single colonies of the two strains were picked and purified twice on Marine Agar 2216 , then inoculated into 3 mL Marine Broth 2216 and incubated at 28 °C with shaking at 200 rpm for 24 h. Cells were harvested by centrifugation at 8,000 × g for 5 min, washed twice with 1× phosphate-buffered saline (1 × PBS; pH 7.4), and resuspended in 1 mL 1× PBS.

The FISH protocol was modified from previous research (*56, 57*). To eliminate artifacts from technical variability, all samples were processed simultaneously on the same slide through identical experimental conditions with matched microscope settings (laser power, gain, and detection range) applied uniformly across all fields of view. A poly-L-lysine-coated glass slide (CITOTEST) was spotted with 20 μL droplets of MC10-B4 suspension (position 1), MC10-B4 suspension (position 2), and MC15-BG7 suspension (position 3). The slide was air-dried in a 37 °C incubator for 30 min, then fixed with 4% paraformaldehyde (Santa Cruz Biotechnology, USA) at 4 °C for 2 h and rinsed twice with 1 × PBS. For permeabilization, samples were treated with a mixture of lysozyme (10 mg/mL) and proteinase K (10 μg/mL) in 0.1 M Tris-HCl (pH 8.0) at 37 °C for 20 min, followed by 1 × PBS rinses twice and sequential dehydration in 50%, 80%, and 100% ethanol (5 min each). The slide was pre-hybridized with a solution containing 5 M NaCl, 1 M Tris-HCl (pH 7.5), 40% formamide, and 10% SDS (0.22 μm-filtered) at 73 °C for 8 min. Hybridization was performed at 42 °C in a humidified chamber overnight. The MC10-specific *parC* probe was then applied to positions 1 and 3, while position 2 received only pre-hybridization solution, serving as blank control. Post-hybridization washes were conducted sequentially in solutions including 50% formamide/2× Saline Sodium Citrate (SSC, 53 °C, 5 min ×2), 0.01% SDS/2× SSC (42 °C, 5 min ×2), 0.5× SSC (42 °C, 5 min ×1), and 0.2× SSC (42 °C, 5 min×1). Samples were counterstained with DAPI (1 μg/mL in PBS, 10 min in darkness), rinsed three times with 1 × PBS, air-dried, and mounted with ProLong Gold anti-fade reagent (Thermo Fisher, USA) under coverslips. Confocal imaging was performed on a Leica Stellaris 8 STED microscope (Leica Microsystems, Germany) using two channels: DAPI (405 nm excitation, 458–547 nm emission) for dsDNA and Cy3 (552 nm excitation, 557–629 nm emission) for the probe signal.

##### 4.6 Tracking the fate of *Ruegeria* MC10-B4 in inoculated corals using FISH

Coral samples were collected before outplanting (**t0**), after 6 and 8 months post-outplanting (**t6** and **t8,** respectively) and fixed in 4% paraformaldehyde at 4 °C overnight for FISH analysis.

FISH procedures were performed following established protocols with modifications (*58, 59*). Fixed samples were decalcified in 20% EDTA solution. After decalcification, samples were washed through a series of ethanol concentrations (50%, 75%, 95%, and 100%), with each step lasting for 15 min. After ethanol dehydration, samples were rinsed twice with xylene, and infiltrated with molten paraffin three times before embedding. Paraffin-embedded specimens were cooled at -20 °C, sectioned into 3 μm slices, mounted on glass slides, and dried at 40 °C overnight. Sections were dewaxed in xylene, washed through an ethanol series (50%, 60%, 70%, 80%, 90%, 100%), and air-dried in the dark.

For probe hybridization, sections were pre-hybridized with 20 μL of pre-hybridization solution (*59*) and incubated for 30 min at 37 °C. A 40 μL of Cy3-labelled probe (1:39 dilution in the pre-hybridization solution) was applied to the samples. Hybridization consisted of a denaturation at 73 °C for 5 min followed by overnight incubation at 42 °C. After hybridization, sections were washed sequentially at predetermined temperatures: 50% formamide/2× SSC 50% formamide/2×SSC at 53 °C for 5 min twice; 0.1% NP-40/2×SSC at 42 °C for 5 min twice; 0.5×SSC and 0.2×SSC at 42 °C for 5 min each. DNA was stained with 20 μL of DAPI for 10 min in the dark, washed with 1x PBS twice, and air-dried. Finally, 20 μL of ProLong Gold anti-fade mounting medium (Thermo Fisher, USA) was applied to the slides to preserve the fluorescence for imaging.

Confocal imaging was performed using a confocal laser scanning microscopy Leica Stellaris 8 STED (Leica Microsystems, Germany) under three channels, i.e., DAPI (excitation: 405 nm, emission: 458–547 nm) for dsDNA, Cy3 (excitation: 552 nm, emission: 557–629 nm) for MC10, and chlorophyll b (excitation: 453 nm, emission: 629–750 nm) for coral autofluorescence. In this setting, the Cy3 channel for probe signal is not overlapping with the coral autofluorescence.

Additionally, a nonsense probe was used to serve as a negative control for FISH on serial slides of the same sample (*59*). Nonsense probe applied in a serial tissue section is essential to differentiate true signals from false positives that were bound to the coral cellular structures, including sporocysts and granular gland cells that are prone to nonspecific probe binding (*66*). The nonsense probe (5’- GTCAGGACCGGGAACGTAGTCAATAGCGTGTCGTCACGGCGTCCGGCGTTTTGATCAGTGCAGACAGGCATCGCAGGCTCCGAGATGTTATGCGGGGAATGTTGGTCGCCATGCCACCGCGATACCGCTGGCACGTTGGCCAGCAGGTTGGGAACTGCGCCGGCAGCACGACGGCTCGGTCAGCGTGCCGCATAGTTGTCGCGGA-3’) was designed based on the antisense sequence of the MC10 *parC* probe. An additional 1 bp deletion every 10 bp of the antisense sequence was conducted to prevent potential hybridization with mRNA expressed by the antisense strand of MC10 *parC (59)*. The non-sense probe was used for FISH on serial slides of the same sample to confirm the hybridization specificity of MC10 *parC* probe.
