## Supplementary figures for "Evolutionary genomics predicts probiotic persistence in corals"

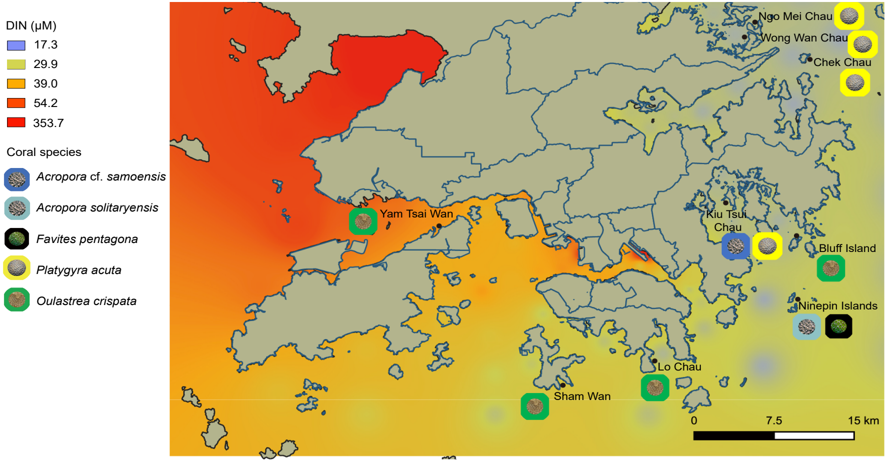


**Fig. S1** A map of Hong Kong showing the concentrations of dissolved inorganic nitrogen (DIN), the sampling sites, and the coral species collected for bacterial isolation. The DIN data is retrieved from the monthly measurement of 76 marine water quality monitoring stations during 1986-2020, published by the Agriculture, Fisheries and Conservation Department (AFCD) of Hong Kong.

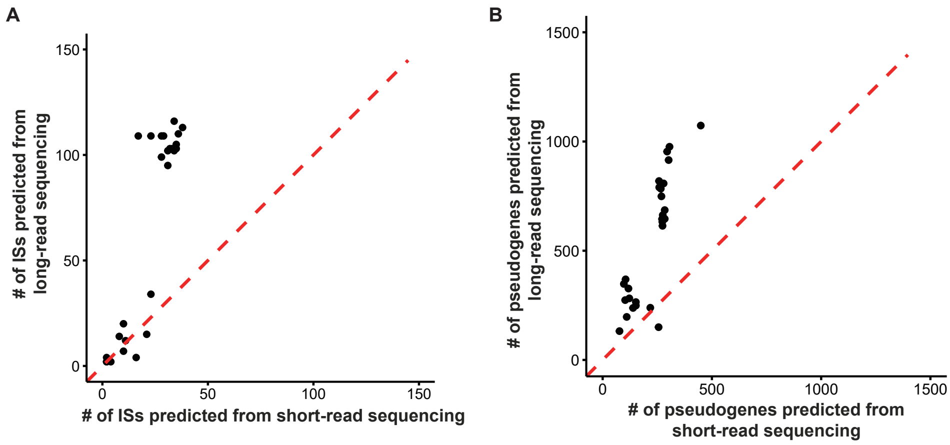


**Fig. S2** Scatter plots comparing (**A**) IS and (**B**) pseudogene counts based on short-read sequencing (DNBSEQ PE150) and long-read sequencing (Nanopore). Each dot represents an isolate sequenced with both platforms. The y = x reference dashed line represents perfect concordance between platforms. Most dots lie above this line, demonstrating systematic underestimation of IS and pseudogene counts by short-read sequencing compared to long-read sequencing.

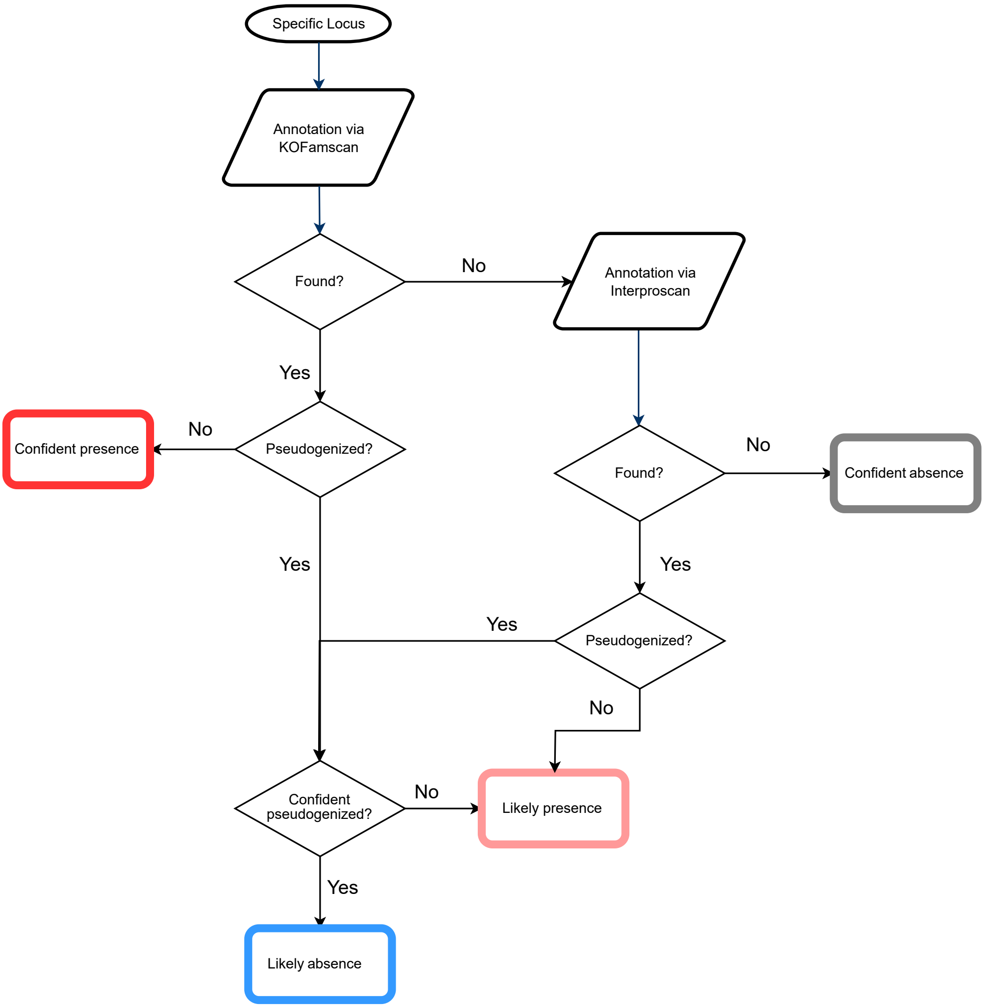


**Fig. S3** The workflow that defines the presence and absence of the gene in a specific pathway as "confident" or "likely". Annotation was first performed using KofamScan, which directly searches gene-based KEGG HMM profiles. To improve functional interpretation, InterProScan was used as a complementary method, as it integrates 13 member databases and covers broader protein domain annotations. Pseudogenes were identified using an adjusted version of Pseudofinder. The confidence of pseudogenization was further assessed based on the intactness of the upstream coding region. A gene was assigned as "Confident presence" if it was successfully annotated by either KOFamScan or InterProScan and showed no evidence of pseudogenization. If a gene was pseudogenized but retained >50% of its upstream region intact, it was categorized as "Likely presence". In cases where a pseudogenized gene had <50% of its upstream region remaining, it was classified as "Likely absence". Finally, if no annotation was detected by KOFamScan or InterProScan, the gene was designated as "Confident absence". The colors of the four brackets are the same as in Fig. 3.

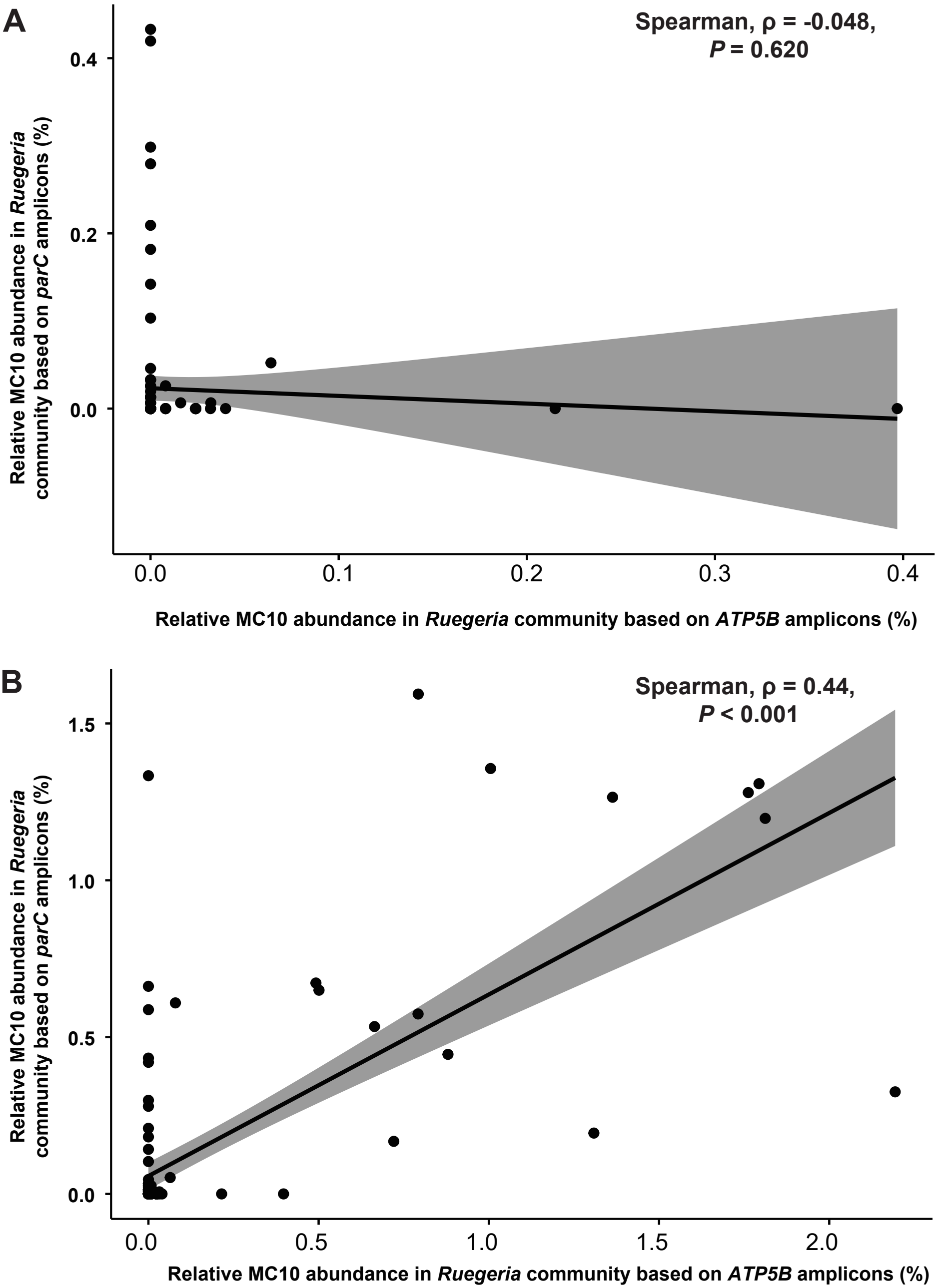


**Fig. S4** Correlation analysis of the relative MC10 abundance in the *Ruegeria* community based on *ATP5B* and *parC* amplicons across two datasets. (**A**) Weak significant correlation was observed between the two marker gene amplicons when relative MC10 abundance is below 0.5%. (**B**) Moderate correlation was observed between the two marker gene amplicons when relative MC10 abundance is between 0% and 2%. Spearman's correlation is considered weak ($|\rho| < 0.3$), moderate ($0.3 \leq |\rho| < 0.5$), and strong ($|\rho| \leq 0.5$).

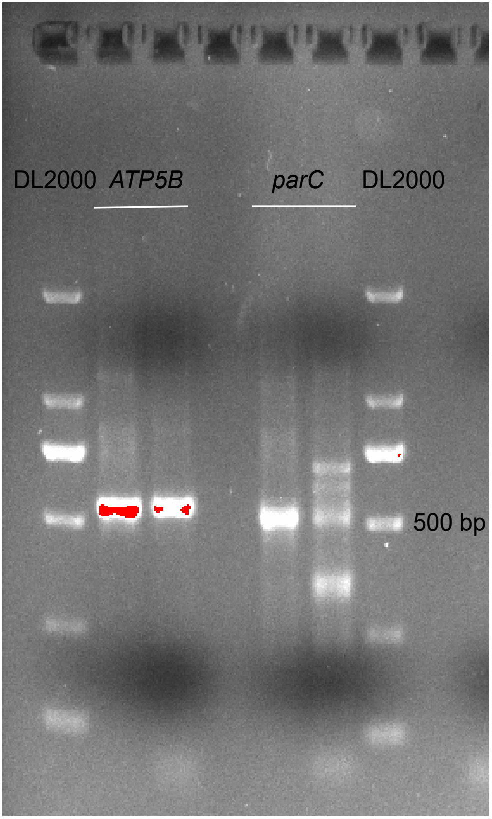


**Fig. S5** Gel electrophoresis analysis of PCR products amplified from *Acropora pruinosa* DNA using *ATP5B*-562F/1111R and *parC*-668F/1189R primer pairs (sequences provided in Table S1). Multiple bands were observed in both reactions, indicating potential non-specific amplification. The amplified product (~500 bp) is larger than the optimal size for conventional qPCR assays. DL2000, DNA marker. The red coloration observed during gel imaging indicates regions of high DNA concentration, a result of ethidium bromide intercalation under UV illumination.

**A**

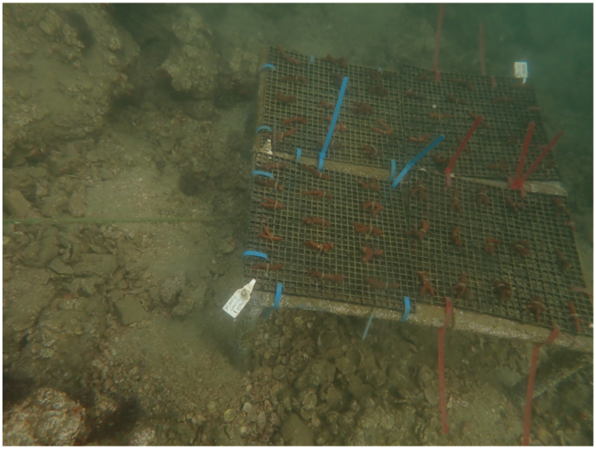

**B**

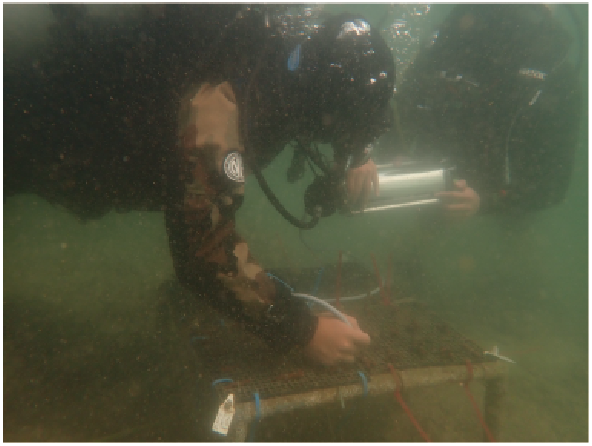

**Fig. S6** (**A**) Overview of the custom-designed coral outplanting platform used in this study. Each platform was comprised of four checkboards, with 15 coral fragments per checkboard (n=60 fragments per platform). (**B**) Divers measuring Fv′/Fm′ for corals underwater using Diving-PAM.

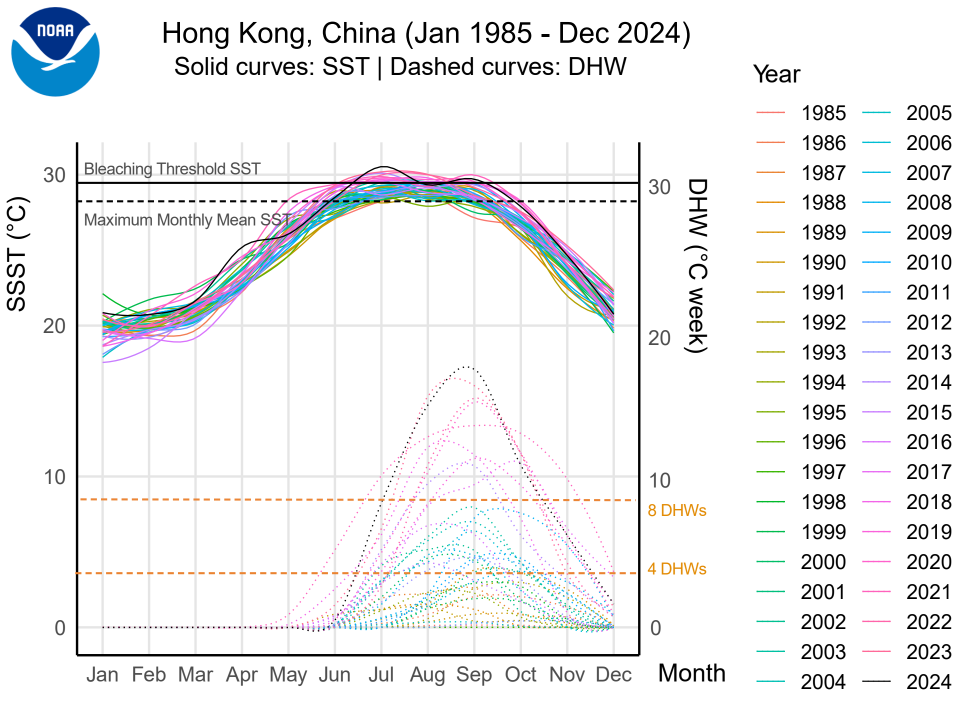


**Fig. S7** Satellite-derived Sea Surface Temperature (SST) shows interannual variability spanning the period from 1985 to 2024 in Hong Kong coastal waters. The Maximum Monthly Mean (MMM) SST indicates the upper boundary of "typical" temperatures, while the Bleaching Threshold SST is set as 1 °C above the MMM. Degree Heating Weeks (DHW in °C weeks, plotted as multiple lines in the bottom right of the panel), defined as the sum of weekly SST anomalies above the MMM threshold, accumulated over 12 consecutive weeks, quantifying cumulative heat stress across the period from 1985 to 2024. All data were obtained from the National Oceanic and Atmospheric Administration (NOAA) Coral Reef Watch products (CRW, V3.1) (60). The 2024 summer thermal event exposed Hong Kong coral communities to severe heat stress, with DHW surpassing 8 °C-weeks.

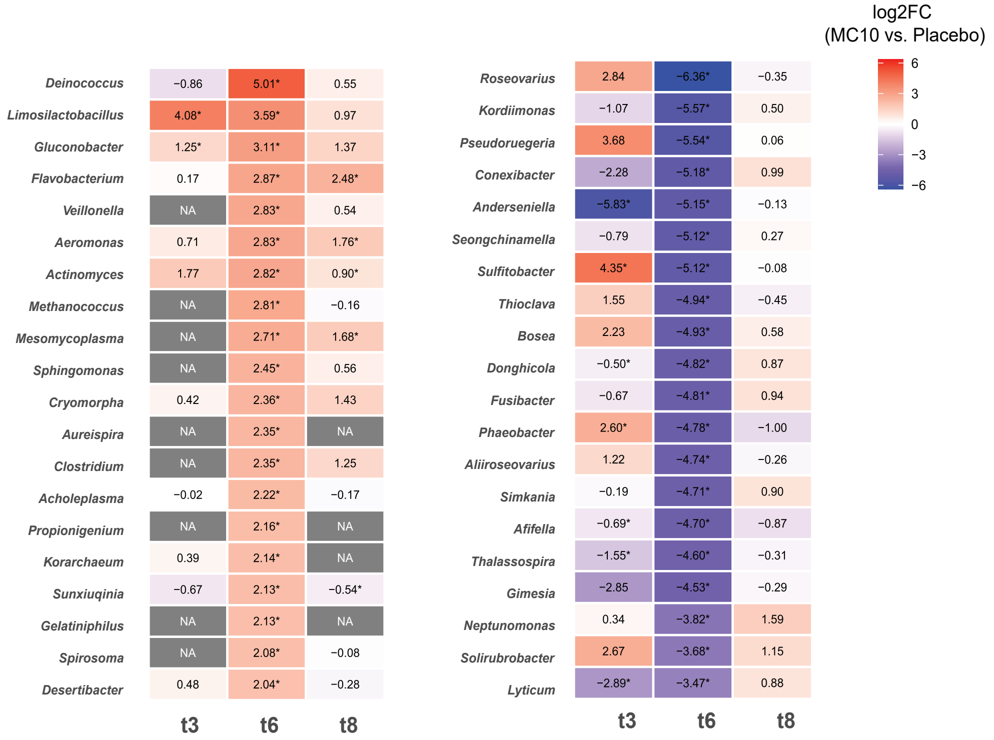


**Fig. S8** The heatmap displays the top 20 significantly upregulated (positive $log_2FC$) and downregulated (negative $log_2FC$) bacterial genera during the bleaching period (t6), mirroring the presentation in Fig. 4B. The visualization further tracks these genera at pre-bleaching (t3) and post-bleaching (t8) phases comparing treatment effects. Missing data values appear as gray tiles, while numeric labels indicate the $log_2$ fold change values (rounded to two decimal places). Asterisks (*) highlight statistically significant differences (FDR-adjusted $P < 0.05$) between treatment groups at each timepoint.

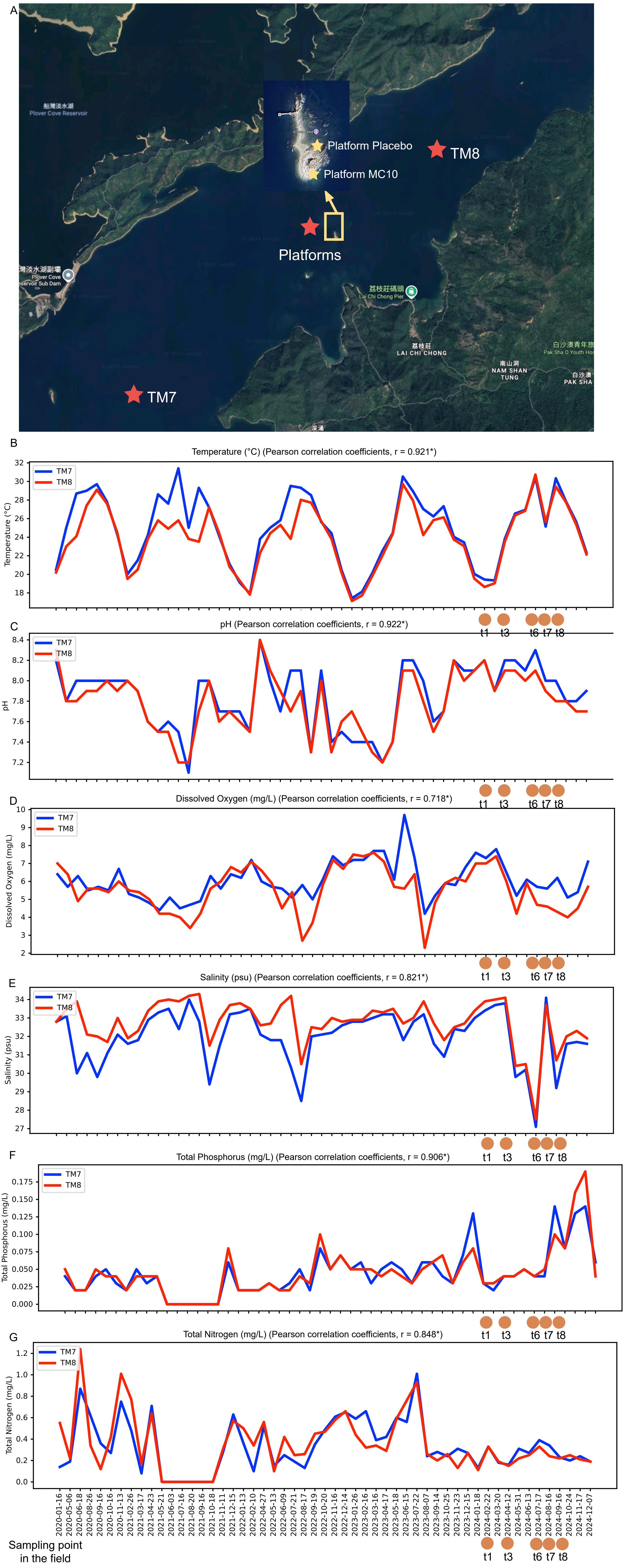


**Fig. S9** Baseline monitoring of abiotic water quality parameters across two outplant sites. (A) Satellite map indicating the locations of experimental platforms (MC10 and Placebo) and environmental monitoring sites TM7 and TM8 within the study area. (B–G) Temporal profiles of key abiotic parameters measured over January 2020 to December 2024 at sites TM7 (blue line) and TM8 (red line). Parameters include (B) Temperature (°C), (C) pH, (D) Dissolved Oxygen (mg/L), (E) Salinity (psu), (F) Total Phosphorus (mg/L), and (G) Total Nitrogen (mg/L). Pearson correlation coefficients indicated high homogeneity between the two sites for all water quality parameters. Orange circles along the x-axis denote the sampling time points during outplant experiment.
